# CAIX forms a transport metabolon with monocarboxylate transporters in human breast cancer cells

**DOI:** 10.1101/625673

**Authors:** Samantha Ames, Jacob T. Andring, Robert McKenna, Holger M. Becker

**Author notes:** Correspondence to: Holger M. Becker.

## Abstract

Tumor cells rely on glycolysis to meet their elevated demand for energy. Thereby they produce significant amounts of lactate and protons, which are exported via monocarboxylate transporters (MCTs), supporting the formation of an acidic microenvironment. The present study demonstrates that carbonic anhydrase IX (CAIX), one of the major acid/base regulators in cancer cells, forms a protein complex with MCT1 and MCT4 in tissue samples from human breast cancer patients, but not healthy breast tissue. Formation of this transport metabolon requires binding of CAIX to the Ig1 domain of the MCT1/4 chaperon CD147 and is required for CAIX-mediated facilitation of MCT1/4 activity. Application of an antibody, directed against the CD147-Ig1 domain, displaces CAIX from the transporter and suppresses CAIX-mediated facilitation of proton-coupled lactate transport. In cancer cells, this “metabolon disruption” results in a decrease in lactate transport, reduced glycolysis and ultimately reduced cell proliferation. Taken together, the study shows that carbonic anhydrases form transport metabolons with acid/base transporters in human tumor tissue and that these interactions can be exploited to interfere with tumor metabolism and proliferation.

## Introduction

Despite substantial progress in early detection and tumor therapy, breast cancer is still the most commonly diagnosed form of cancer and leading cause of cancer death in women worldwide [1]. Like most solid tumors, breast tumor tissue displays a significant increase in glycolytic activity, as compared to healthy breast tissue [2]). This increase in glycolysis, which was discovered by Otto Warburg more than 90 years ago [3, 4], is now considered an emerging hallmark of cancer [5]. Glycolytic activity results in the production of lactate and protons, which have to be removed from the cell to avoid intracellular acidosis and suffocation of metabolism. Lactate flux in cancer cells is primarily mediated by the monocarboxylate transporters MCT1 and MCT4, both of which transport lactate together with H^+^ in an electroneutral 1:1 stoichiometry [6–9]. MCT1, which is ubiquitously expressed in most tissue, has a K_m_ value of 3-5 mM for lactate [6, 7] and mediates both influx and efflux of lactate and protons, depending on the cell’s metabolic profile. MCT4 is considered a high-capacity carrier with a K_m_ value of 20-35 mM for lactate [8], which primarily acts as a lactate exporter in glycolytic cells, including hypoxic cancer cells. Expression of both MCT1 and MCT4 is increased in breast cancer tissue, as compared to healthy breast epithelium, and correlates with increasing tumor grade [10–15]. Furthermore, expression of MCT1 was shown to correlate with shorter progression-free survival and increased risk of recurrence after chemotherapy in breast cancer patients [2, 10]. In cultured breast cancer cells, knockdown of MCT1 and MCT4, decreases proliferation, migration and invasion [16]. These finding indicate that MCT-mediated lactate/H^+^ cotransport contributes to cancer cell aggressiveness.

Trafficking of the MCT proteins to the plasma membrane, but also regulation of transport function, is mediated by ancillary proteins. For MCT1 and MCT4 the preferred ancillary protein is CD147 which is tightly associated with the transporter in the membrane [17–19]. The general isoform of CD147 (termed CD147-Ig1-Ig2, basigin or basigin-1), which is ubiquitously expressed in most tissue, comprises two extracellular, immunoglobulin (Ig)-like domains, a transmembrane domain and a short intracellular C-terminal tail [20, 21]. CD147 is a multifunctional protein, which has been attributed a key role in the development and progression of different tumor types, including breast cancer. CD147 was shown to induce surface expression of matrix metalloproteases in adjacent tumor cells or fibroblast and has therefore been attributed a function in tumor cell invasion [22, 23]. Furthermore, CD147 can induce expression of vascular endothelial growth factors (VEGF) and production of hyaluronan to stimulate angiogenesis [22]. CD147 interacts with the multidrug resistance transporter MDR1 and vacuolar H^+^-ATPase, thereby promoting chemoresistance [24, 25]. Immunohistological staining demonstrated that expression of CD147 is upregulated in breast cancer tissue, as compared to normal breast tissue [10]. Similar to the expression of MCT1 and MCT4, expression levels of CD147 increase with increasing histological grade [10, 15] and positively correlate with poor prognosis and chemoresistance in breast cancer patients [10, 23, 25].

Transport activity of MCTs can be facilitated by carbonic anhydrase IX (CAIX), an enzyme that has been attributed a central role in tumor acid/base regulation and cancer progression [26–30]. CAIX is tethered to the extracellular face of the plasma membrane via a transmembrane domain. Like the other isoforms of carbonic anhydrase, catalytic activity of CAIX is supported by an intramolecular proton shuttle, which mediates the rapid exchange of H^+^ between the enzyme’s catalytic center and the surrounding bulk solution. Unlike the other carbonic anhydrases however, CAIX features a 59 amino acid long proteoglycan-like (PG) domain, which is unique to CAIX within the CA family [28].

In healthy tissue, expression of CAIX, which is under control of the hypoxia-inducible factor HIF-1α [31], is restricted to intestine and gall bladder [32]. However, CAIX is upregulated in many different tumor types, including breast cancer [33, 34]. Expression levels of CAIX positively correlate with histological tumor grade in breast cancer [12–14, 34–40]. Furthermore, expression of CAIX was found to correlate with reduced overall survival and higher occurrence of relapse and is associated with chemoresistance in breast cancer patients [34, 41, 42]. Interestingly, expression of CAIX was also found to correlate with MCT1 and CD147 in breast tumor tissue [34].

CAIX physically and functionally interacts with various acid/base transporters in tumor cells. CAIX was shown to colocalize with the Na^+^/HCO_3_^−^ cotransporter NBCe1 and the Cl^−^ /HCO_3_^−^ exchanger AE2 in the lamellipodia of migrating SiHa cells [43]. Therefore it was suggested that CAIX can form a “transport metabolon” with NBCe1 and AE2 to facilitate HCO_3_^−^ flux across the cell membrane and drive cell migration [43]. A “transport metabolon” is defined as a structural and functional complex, composed of a transport protein and an enzyme, which interact with each other to facilitate substrate transport across the cell membrane. Transport metabolons are formed from various intracellular and extracellular carbonic anhydrases and acid/base transporters, including Na^+^/H^+^-exchangers (NHEs), AEs, NBCs, and MCTs (for review see [44–47]. We have previously shown that CAIX facilitates MCT transport activity in cultured breast cancer cells and when heterologously coexpressed in *Xenopus* oocytes, by a mechanism that is independent from the CA’s catalytic function [29, 30]. In the present study, we investigate whether CAIX forms a transport metabolon with MCT1 and MCT4 in human breast cancer tissue and analyze the structural components that mediate the formation of this protein complex. Furthermore, we elaborate whether the MCT1/4-CAIX transport metabolon can be exploited as drug target for tumor therapy.

## Results

### CAIX interacts with MCT1 and MCT4 in human breast cancer cells

Lactate efflux from cancer cells is primarily mediated by the monocarboxylate transporters MCT1 and MCT4 [10, 11, 48], the activity of which has been shown to be facilitated by CAIX, which functions as proton antenna for the transporter [29, 30]. In order to establish an efficient proton transfer, enzyme and transporter have to be localized in close proximity to each other. To investigate whether CAIX forms such a “transport metabolon” with MCTs in human breast cancer tissue, we first analyzed the physical interaction between CAIX and MCT1/MCT4, using *in situ* proximity ligation assays (PLA). CAIX was found to interact with MCT1 in tumor tissue samples from human breast cancer patients (Figure 1 A-C, E), but not in uninvolved breast tissue (Figure 1, D, E). Interestingly, the number of protein complexes increased in samples from grade II and grade III tumors, as compared to grade I tumors. CAIX did also interact with MCT4 in the same set of tissue samples (Figure 2 A-C, E), but not in uninvolved breast tissue (Figure 2 D, E). While no significant changes in the number of protein complexes could be detected between grade I and grade II tumors, grade III tumors showed a significant increase in the number of MCT4-CAIX interactions. When MCT1 and MCT4 are considered together, the number of transport metabolons constantly increases with increasing tumor grade from 13 signals/cell in grade I, 20 signals/cell in grade II and 30 signals/cell in grade III tumors.

**Figure 1:**
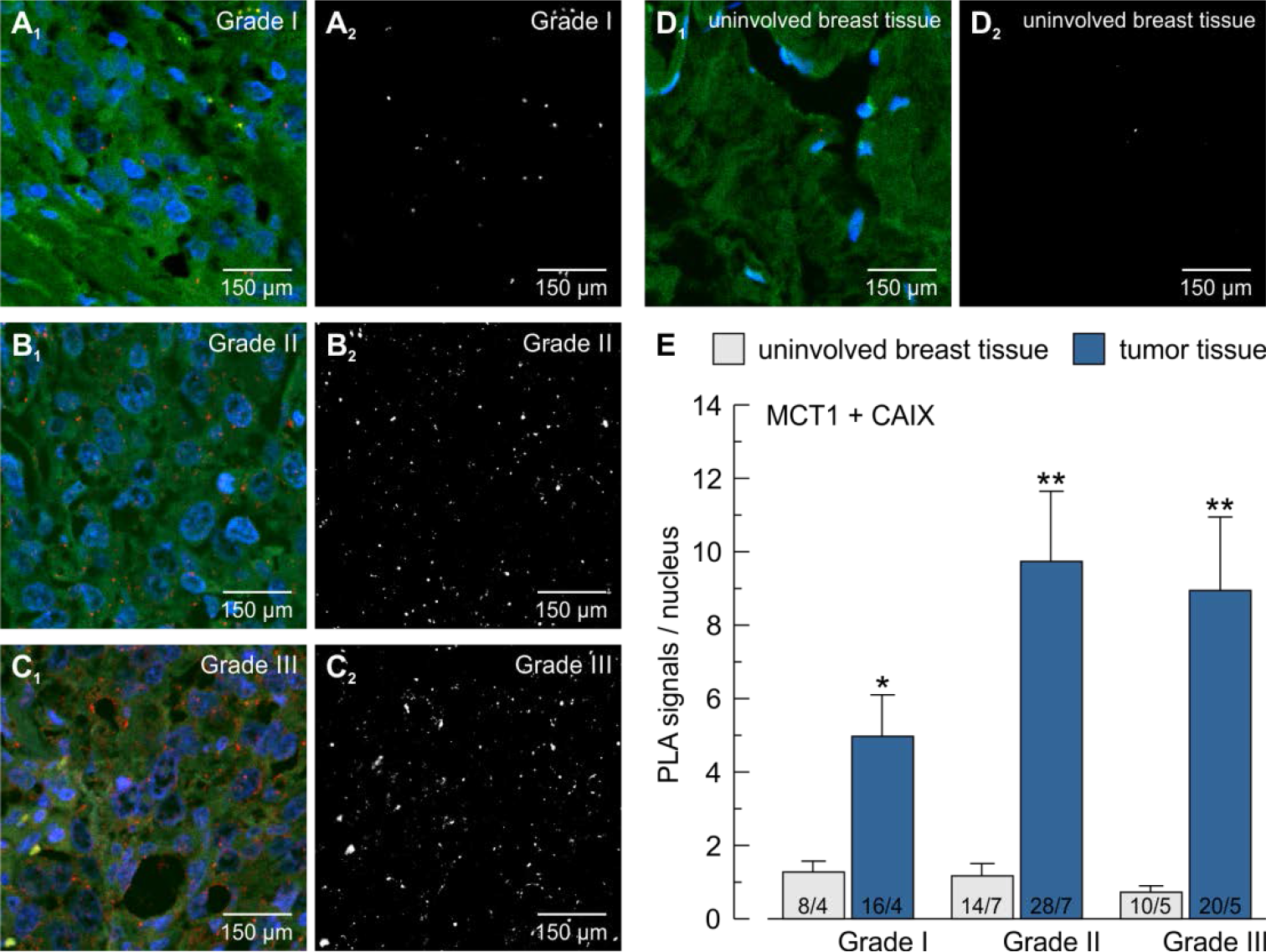
CAIX is colocalized with MCT1 in human breast tumor tissue, but not in healthy breast tissue. (**A**-**D**) *In situ* proximity ligation assay (PLA) for CAIX and MCT1 in grade I (**A**), grade II (**B**), and grade III (**C**) breast tumor tissue, and uninvolved breast tissue (**D**) of human cancer patients. Panels A_1_, B_1_, C_1_, and D_1_ show the PLA signals (red), nuclei staining (blue) and actin staining (green). Panels A_2_, B_2_, C_2_, and D_2_ show exclusively the PLA signals. (**E**) Quantification of the PLA signals per nucleus. The asterisks above the bars for tumor tissue refer to the corresponding bars for uninvolved breast tissue.

**Figure 2:**
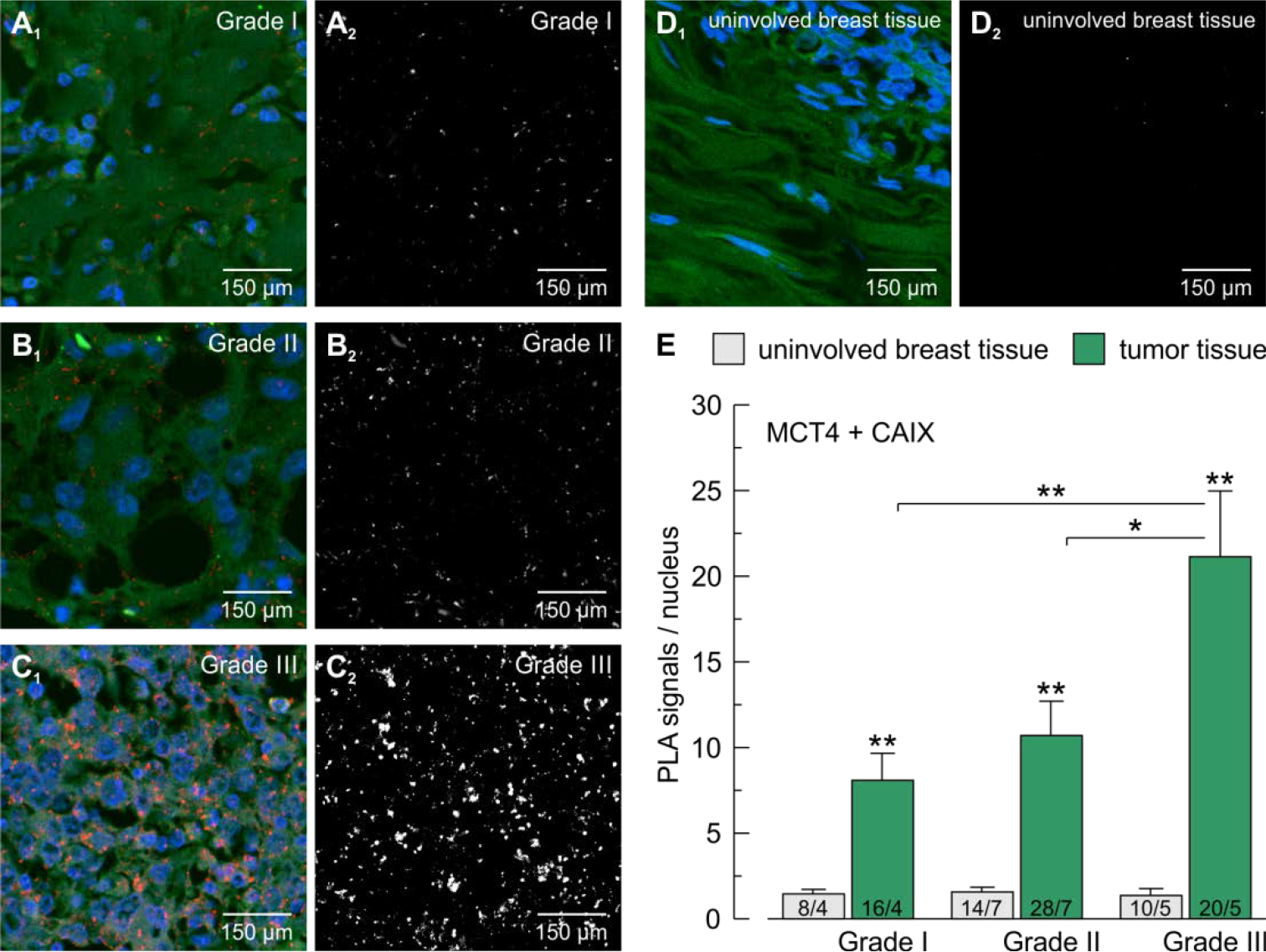
CAIX is colocalized with MCT4 in human breast tumor tissue, but not in healthy breast tissue. (**A**-**D**) *In situ* proximity ligation assay (PLA) for CAIX and MCT4 in grade I (**A**), grade II (**B**), and grade III (**C**) breast tumor tissue, and uninvolved breast tissue (**D**) of human cancer patients. Panels A_1_, B_1_, C_1_, and D_1_ show the PLA signals (red), nuclei staining (blue) and actin staining (green). Panels A_2_, B_2_, C_2_, and D_2_ show exclusively the PLA signals. (**E**) Quantification of the PLA signals per nucleus. The asterisks above the bars for tumor tissue refer to the corresponding bars for uninvolved breast tissue.

Expression of CAIX is controlled by the hypoxia-inducible factor, HIF-1α. Therefore, it appears likely that the MCT-CAIX transport metabolon will primarily form under hypoxic conditions. Indeed, the number of interactions between MCT1 and CAIX, as well as MCT4 and CAIX more than doubled in MDA-MB-231 breast cancer cells, incubated under hypoxia (1% O_2_), as compared to normoxic cells (Figure 3 A_1_, A_2_, B_1_, B_2_, C, D). In MCF-7 cells, hypoxia induced a more than threefold increase in MCT1-CAIX interactions (Figure 4 A_1_, A_2_, B). In both cell lines, knockdown of CAIX resulted in a significant decrease in the PLA signal, while omission of the primary antibodies resulted in no signal at all (Figure 3 A_3_, A_4_, B_3_, B_4_, C, D; Figure 4 A_3_, A_4_, B). In MCF-7 cells, which do not express MCT4 [29], no signals where observed when the assay was carried out with antibodies against MCT4 and CAIX (Figure 4 B, right bar). Colocalization of MCT1/4 and CAIX could also be confirmed by conventional immune staining for MCT1, MCT4 and CAIX in MDA-MB-231 (Figure 3 E_1_, E_2_) and MCF-7 cells (Figure 4 C_1_, C_2_, C_3_, C_4_). Taken together, these results indicate that MCT1 and MCT4 can form a protein complex with CAIX in hypoxic breast cancer cells.

**Figure 3:**
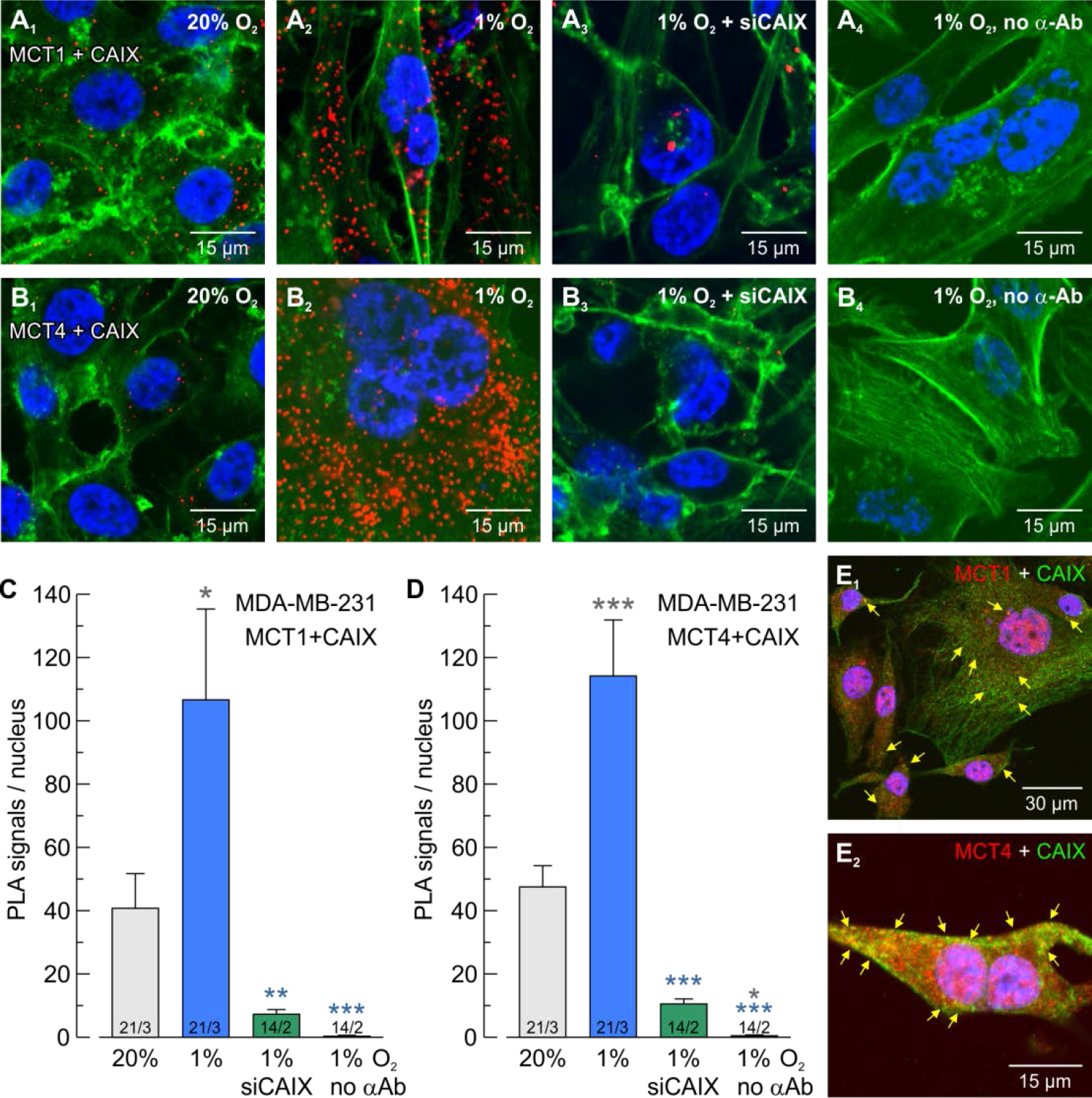
CAIX is colocalized with MCT1 and MCT4 in hypoxic MDA-MB-231 breast cancer cells. (**A**, **B**) *In situ* proximity ligation assay (PLA) for CAIX + MCT1 (**A**) and CAIX + MCT4 (**B**), respectively, in normoxic (**A**_**1**_, **B**_**1**_) and hypoxic (**A**_**2**_, **B**_**2**_) MDA-MB-231 cells. For control CAIX was knocked down with siRNA (**A**_**3**_, **A**_**4**_) or the PLA was performed without primary antibodies (**A**_**4**_, **B**_**4**_). PLA signals are shown in red, nuclei in blue and actin in green. (**C**, **D**) Quantification of the PLA signals for CAIX + MCT1 (**C**) and CAIX + MCT4 (**D**), respectively. The asterisks of a given color refer to the bars of the corresponding color. (**E**) Antibody staining for CAIX (green) and MCT1 (**E**_**1**_, red) and MCT4 (**E**_**2**_, red), respectively, in hypoxic MDA-MB-231 cells. Nuclei are shown in blue.

**Figure 4:**
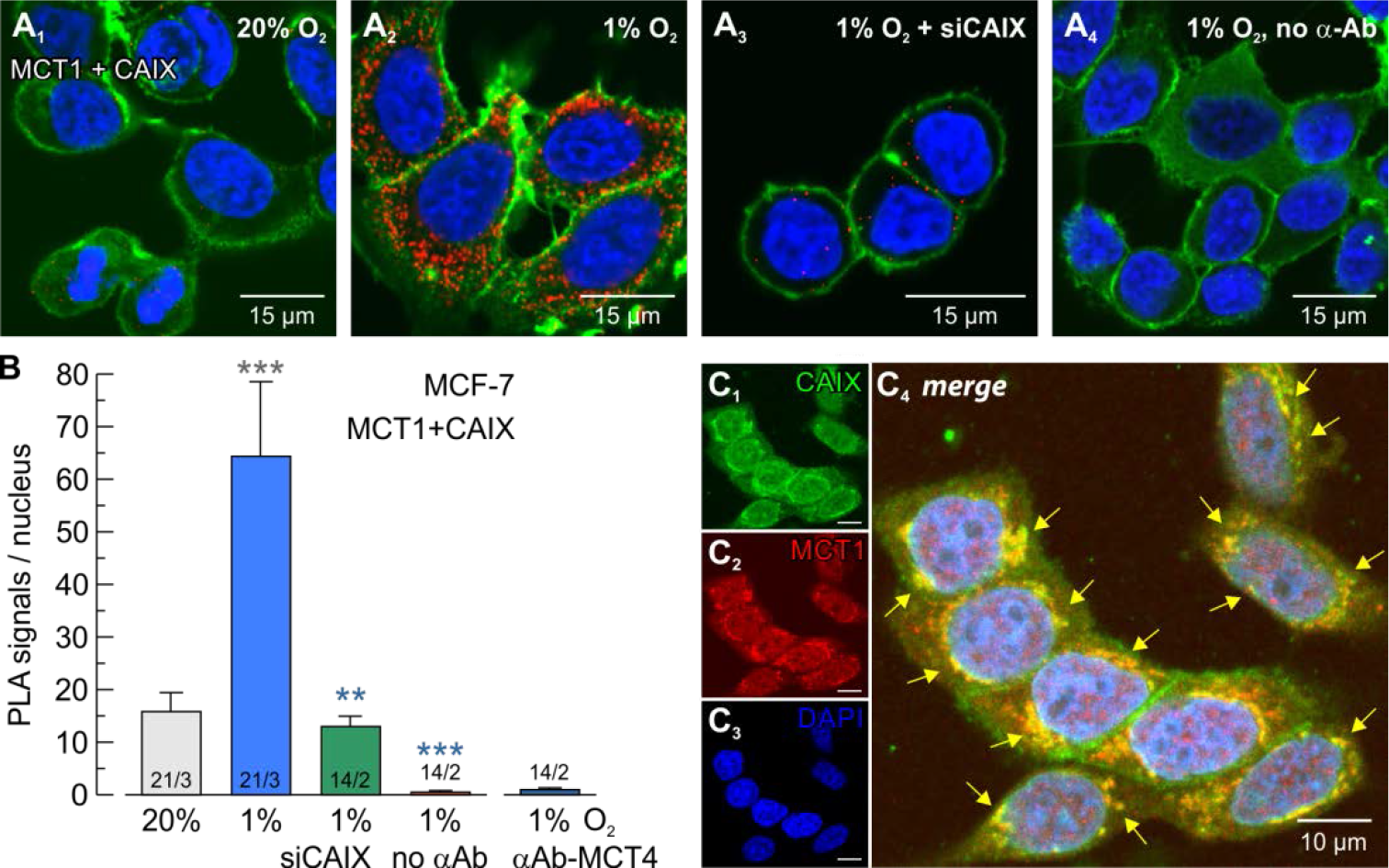
CAIX is colocalized with MCT1 in hypoxic MCF-7 breast cancer cells. (**A**) *In situ* proximity ligation assay (PLA) for CAIX and MCT1 in normoxic (**A**_**1**_) and hypoxic (**A**_**2**_) MCF-7 cells. For control CAIX was knocked down with siRNA (**A**_**3**_) or the PLA was performed without primary antibodies (**A**_**4**_). PLA signals are shown in red, nuclei in blue and actin in green. (**B**) Quantification of the PLA signals for CAIX and MCT1. The asterisks of a given color refer to the bars of the corresponding color. (**C**) Antibody staining for CAIX (**C**_**1**_, green), MCT1 (**C**_**2**_, red) and nuclei (**C**_**3**_, blue). **C**_**4**_: Overlay of signals shown in C_1-3_.

### CAIX binds to the Ig1 domain of the MCT1/4 chaperon CD147

Direct interaction between MCT1/4 and CAIX requires binding between the proteins. We have recently shown that the extracellular CA isoform CAIV does not directly bind to MCT1 or MCT4, but to the Ig1 domain in the transporters’ chaperon CD147 [49]. Binding of CAIX to CD147 was tested by pulldown of CAIX, heterologously expressed in *Xenopus* oocytes, with a GST fusion protein of the Ig1 domain of CD147 (Figure 5). Pulldown of CAIX with CD147-WT resulted in a robust signal for CAIX, indicating direct binding between the two proteins (Figure 5 C, D). Molecular docking experiments suggested that binding between CD147 and CAIX is mediated by CD147-Glu73 (Figure 5 A_1_) and CAIX-His200 (Figure 5 A_2_). Indeed, mutation of Glu73 in the Ig1 domain of CD147 and His200 in the catalytic domain of CAIX resulted in a loss of interaction, while mutation of the adjacent Glu31 in the Ig1 domain of CD147 had no significant effect on binding (Figure 5 C, D). Equal expression of CAIX-WT and CAIX-H200A in *Xenopus* oocytes, used for the pulldown, was confirmed by western blot analysis (Figure 5 B). In summary, the pulldown shows that CAIX binds to the Ig1 domain of CD147, most likely by forming a hydrogen bond between CAIX-His200 and CD147-Glu73.

**Figure 5:**
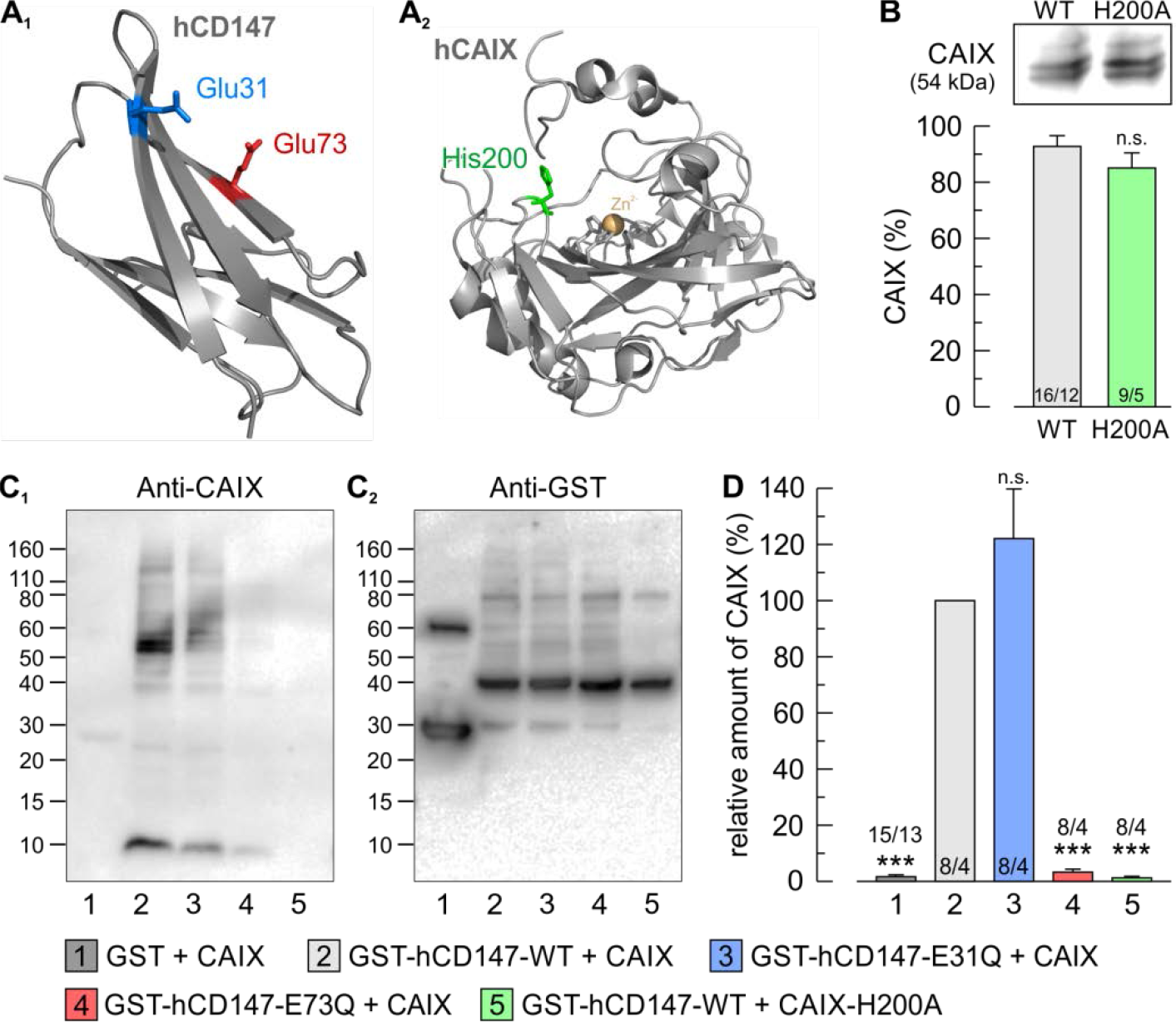
CAIX binds to the Ig1 domain of the MCT1/4 chaperon CD147. (**A**) Structure of the Ig1 domain of (**A**_**1**_) human CD147 (PDB-ID: 3B5H [89]) and (**A**_**2**_) human CAIX (PDB-ID: 4ZAO [90]). The putative binding sites (hCD147-E31, hCD147-E73 and CAIX-H200) are color labelled. (**B**) Western blot against CAIX from oocytes expressing CAIX-WT or CAIX-H200A. The antibody recognizes both WT and mutant equally well. (**C**) Representative western blots for CAIX (**C**_**1**_) and GST (**C**_**2**_). CAIX-WT and CAIX-H200A, respectively, was pulled down with a GST fusion protein of the Ig1 domain of hCD147-WT or a mutant of the protein. (**D**) Relative fluorescent signal of CAIX. For every blot, the signals for CAIX were normalized to the corresponding signals for GST-hCD147-WT. Each individual signal for CAIX was normalized to the intensity of the signal for GST in the same lane. Mutation of CAIX-H200 and hCD147-E73, but not hCD147-E31, abolishes binding between CAIX and hCD147.

### CAIX-mediated facilitation of MCT transport activity requires binding of CAIX to CD147

To investigate whether direct binding of CAIX to CD147 is required for CAIX-mediated facilitation of MCT transport activity, we coexpressed rat MCT1 together with rat CD147-WT or a mutant of the chaperon and CAIX in *Xenopus* oocytes (Figure 6 A). Coexpression of MCT1 and rCD147-WT with CAIX resulted in a significant increase in MCT1 transport activity, both in the direction of influx and efflux, as measured from the rate of change in intracellular H^+^ concentration during application (Figure 6 C) and removal (Figure 6 D) of lactate. Mutation of rCD147-Lys73 (the analogue to hCD147-Glu73) resulted in a loss of functional interaction between MCT1 and CAIX, while mutation of rCD147-Glu32 (analogue to hCD147-Glu31) had no effect on CAIX-mediated facilitation in MCT1 activity (Figure 6 A, C, D). Neither mutation of rCD147-Lys73 nor rCD147-Glu32 had any effect on MCT1 transport activity in the absence of CAIX (Figure 6 C, D). Expression of CAIX was confirmed by a CO_2_ pulse at the end of each experiment (Figure 6 A, B). Extracellularly applied CO_2_ diffuses into the oocyte, where it is hydrated to HCO_3_^−^ and H^+^, resulting in intracellular acidification. Since hydration of CO_2_ is catalyzed by CAIX, the rate of intracellular acidification provides a direct measure for CAIX catalytic activity. That CAIX-His200 is crucial for the functional interaction between MCT1 and CAIX was already confirmed in a previous study (see Figure 4 in [29]).

**Figure 6:**
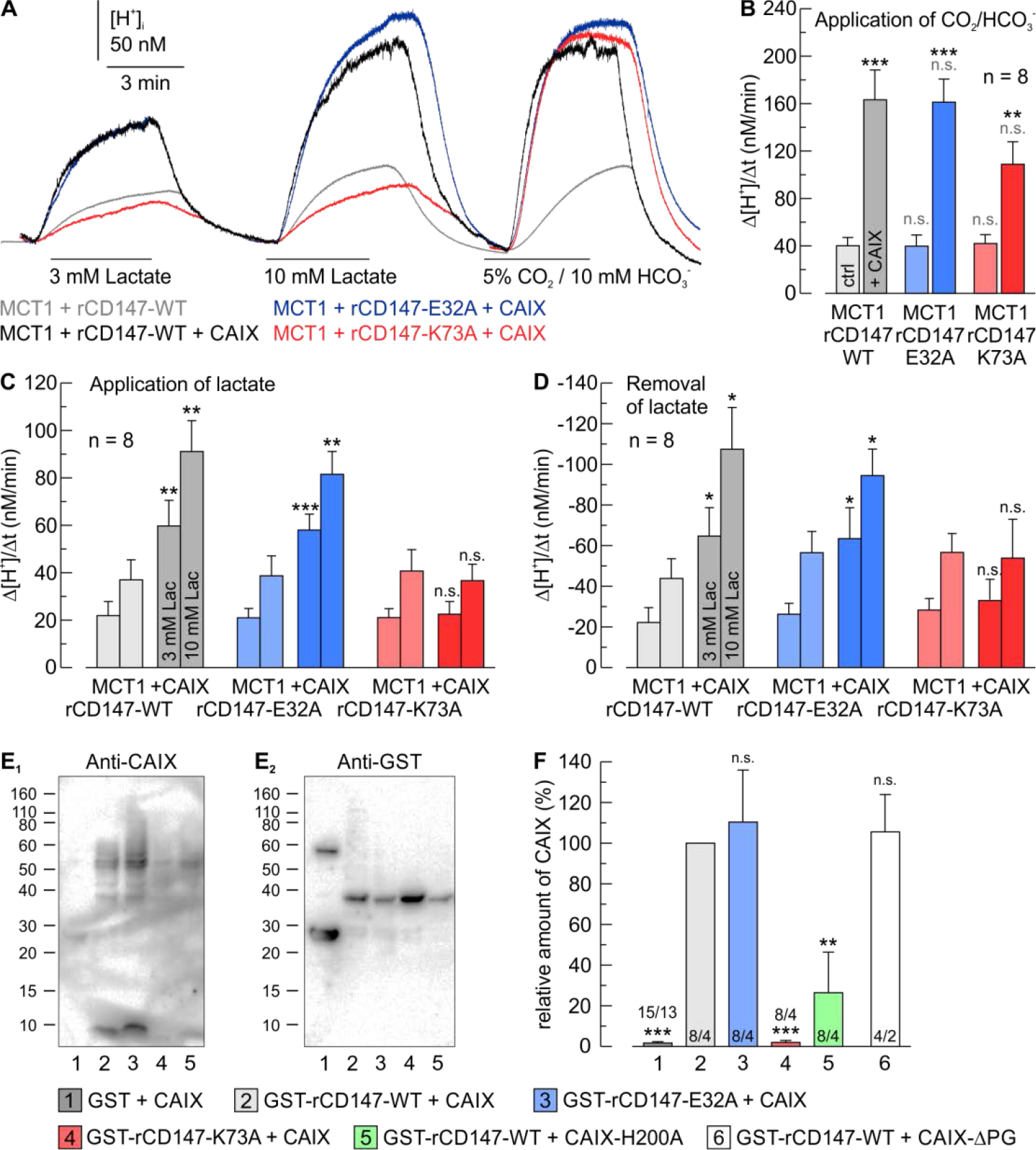
CAIX-mediated facilitation of MCT1 transport activity requires binding of CAIX to the CD147 Ig1 domain. (**A**) Original recordings of the change in intracellular H^+^ concentration during application of lactate and CO_2_/HCO_3_^−^ in *Xenopus* oocytes, expressing rMCT1 together with rCD147-WT (gray traces), rCD147-E32A (blue traces) or rCD147-K73A (red traces), either without (light traces) or with CAIX (dark traces). (**B**-**D**) Rate of change in intracellular H^+^ concentration (Δ[H^+^]/Δt) duringapplication of 5% CO_2_ / 10 mM HCO_3_^−^ (**B**)and application (**C**) and removal of lactate (**D**) in *Xenopus* oocytes, expressing rMCT1 together with rCD147-WT (gray bars), rCD147-E32A (blue bars), or rCD147-K73A (red bars), either without (light bars) or with CAIX (dark bars). The asterisks above the bars for CAIX-expressing cells refer to the corresponding bars for non-CAIX-expressing oocytes. Mutation of rCD147-K73, but not rCD147-E32, abolishes CAIX-mediated facilitation of MCT1 transport activity. (**E**) Representative western blots for CAIX (**E**_**1**_) and GST (**E**_**2**_). CAIX-WT and CAIX-H200A, respectively, was pulled down with a GST fusion protein of the Ig1 domain of hCD147-WT or a mutant of the protein. (**F**) Relative fluorescent signal of CAIX. For every blot, the signals for CAIX were normalized to the corresponding signals for GST-rCD147-WT. Each individual signal for CAIX was normalized to the intensity of the signal for GST in the same lane. Mutation of CAIX-H200 and rCD147-K73, but not rCD147-E32, abolishes binding between CAIX and rCD147. The CAIX-PG domain is not involved in binding to CD147, since the truncation mutant CAIX-ΔPG binds GST-rCD147 equally well as CAIX-WT.

To confirm that binding of CAIX to rCD147 follows the same mechanism as observed for hCD147, CAIX was pulled down with a GST fusion protein of the Ig1 domain of rCD147-WT or a mutant of the protein (Figure 6 E, F). Pulldown of CAIX with rCD147-WT and rCD147-E32A resulted in a robust signal for CAIX, while pulldown with rCD147-K73A produced no signal for CAIX (Figure 6 F). Pulldown of CAIX-H200A with rCD147-WT also resulted in no signal for CAIX in three out of four pulldowns. In one of the four pulldowns, however, a signal for CAIX could be detected for unknown reason (Figure 6 F). We have recently shown that CAIX-mediated facilitation of MCT transport activity involves the CAIX proteoglycan-like (PG) domain, which could function as proton antenna for the protein complex [30]. To investigate whether the PG domain might also play a role in the binding of CAIX to CD147, the CAIX mutant CAIX-ΔPG, missing the PG domain, was pulled down with GST-rCD147 (Figure 6 F, white bar). However, pulldown of CAIX-ΔPG resulted in a similar signal for CAIX as observed for CAIX-WT, indicating that the PG domain is not involved in the binding of CAIX to CD147.

Transport of lactate in cancer cells is primarily mediated by MCT1 and MCT4. However, some cancer cells, including breast carcinoma, colon adenocarcinoma, nonsmall cell lung cancer, ovarian adenocarcinoma and prostate cancer, do also express MCT2 [11,], the chaperon of which is GP70 [18]. The Ig1 domain of GP70 carries a putative CAIX binding site (GP70-Asp95, GP70-Arg130) that is analogue to the binding site in CD147 (Figure 7 A). Both residues are conserved in the rat and human isoforms of GP70. To investigate whether CAIX facilitates transport activity of MCT2 and whether this facilitation requires direct binding of CAIX to the transporter’s chaperon GP70, we coexpressed rat MCT2 (rMCT2) together with rat GP70-WT (rGP70-WT) or a mutant of the protein and CAIX in *Xenopus* oocytes (Figure 7 B). Coexpression of rMCT2 and rGP70-WT with CAIX resulted in a significant increase in MCT2 transport activity, both in the direction of influx and efflux (Figure 7 C, D). Mutation of rGP70-Arg130 resulted in a loss of functional interaction between MCT2 and CAIX, while mutation of rGP70-Asp95 had no effect on CAIX-mediated facilitation in MCT2 activity (Figure 7 B, C, D). Neither mutation of rGP70-Arg130 nor rGP70-Asp95 had an effect on MCT2 transport activity in the absence of CAIX (Figure 7 C, D). Expression of CAIX was confirmed by a CO_2_ pulse at the end of each experiment (Figure 7 A). Pulldown of CAIX with a GST-fusion protein of rGP70-WT resulted in a robust signal for CAIX, indicating direct binding between the two proteins (Figure 7 E, F). Mutation of rGP70-Arg130 and CAIX-His200, respectively, resulted in a loss of interaction, while mutation of rGP70-Asp95 had no significant effect on binding (Figure 7 E, F). These results indicate that CAIX could also form a transport metabolon with MCT2/GP70 to facilitate lactate flux in MCT2-expressing cancer cells.

**Figure 7:**
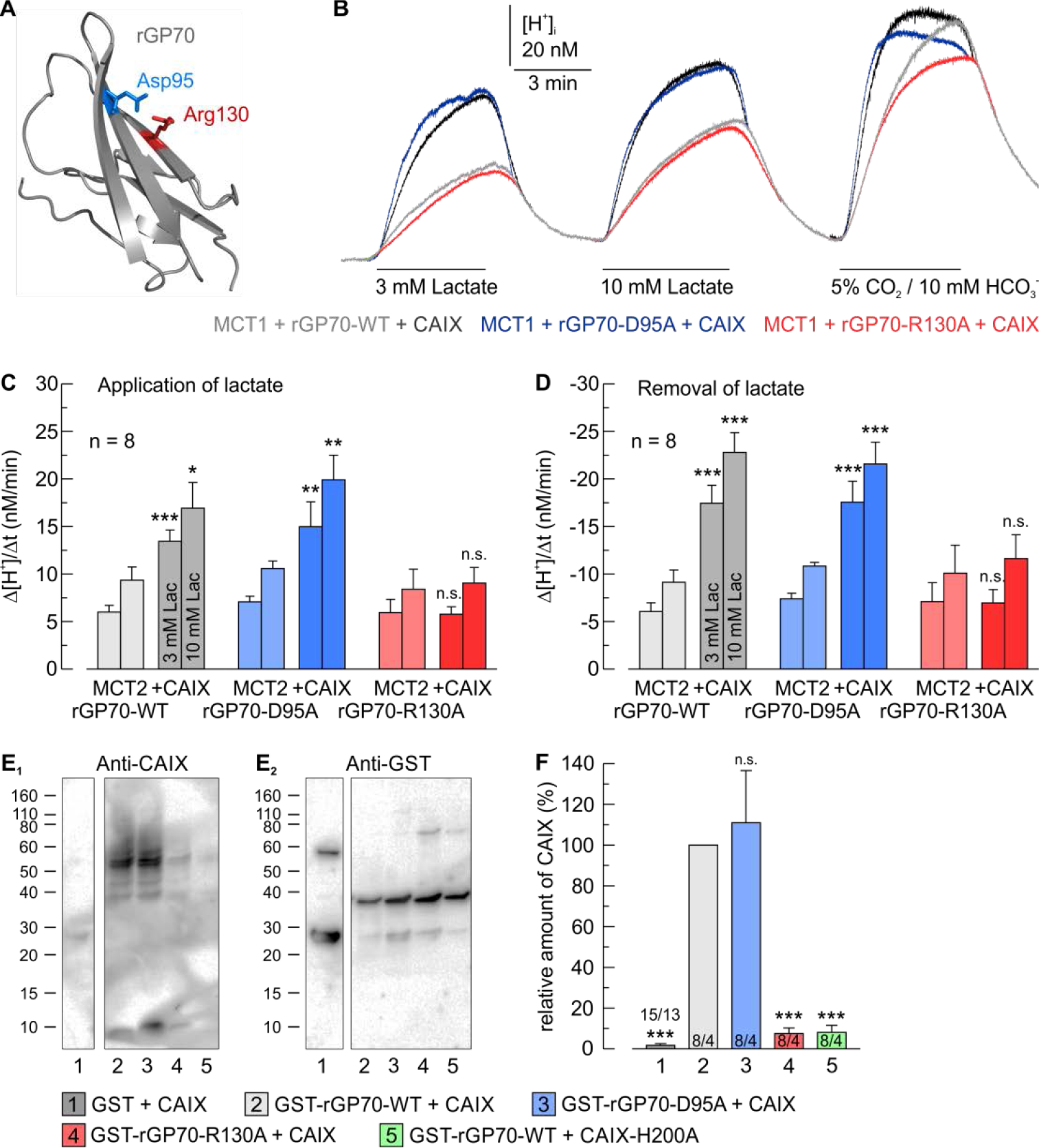
CAIX-mediated facilitation of MCT2 transport activity requires binding of CAIX to the Ig1 domain of the MCT2 chaperon GP70. (**A**) Homology structure of the Ig1 domain of rat GP70 (based on human CD147 (PDB-ID: 3B5H [89])). The putative CAIX binding sites (GP70-D95 and GP70-R130) are color labelled. (**B**) Original recordings of the change in intracellular H^+^ concentration during application of lactate and CO_2_/HCO_3_^−^ in *Xenopus* oocytes, expressing rMCT2 together with rGP70-WT (gray traces), rGP70-D95A (blue traces), or rGP70-R130A (red traces), either without (light traces) or with CAIX (dark traces). (**C**, **D**) Rate of change in intracellular H^+^ concentration (Δ[H^+^]/Δt) during application (**C**) and removal of lactate (**D**) in *Xenopus* oocytes, expressing rMCT2 together with rGP70-WT (gray bars), rGP70-D95A (blue bars), or rGP70R130A (red bars), either without (light bars) or with CAIX (dark bars). The asterisks above the bars for CAIX-expressing cells refer to the corresponding bars for non-CAIX-expressing oocytes. Mutation of rGP70-R130, but not D95, abolishes CAIX-mediated facilitation of MCT2 transport activity. (**E**) Representative western blots for CAIX (upper blot) and GST (lower blot). CAIX-WT and CAIX-H200A, respectively was pulled down with a GST fusion protein of the Ig1 domain of rGP70-WT or a mutant of the protein. (**F**) Relative fluorescent signal of CAIX. For every blot, the signals for CAIX were normalized to the corresponding signals for GST-rGP70-WT. Each individual signal for CAIX was normalized to the intensity of the signal for GST in the same lane. Mutation of CAIX-H200 and rGP70-R130, but not rGP70-D95, abolishes binding between CAIX and rGP70.

### Functional interaction between MCT and CAIX can be disrupted by an antibody against CD147

Inhibition of lactate efflux, either by direct inhibition of MCT transport activity [29, 52, 53] or knockdown of CAIX [29], was shown to significantly decrease breast cancer cell proliferation. Since expression of CAIX is predominately associated with cancer cells, the MCT1/4-CD147-CAIX transport metabolon appears as a promising target to interfere with proton/lactate transport in cancer cells. Therefore, we investigated whether the CAIX-mediated increase in MCT transport activity could be abolished by an interfering agent which disrupts binding of CAIX to the transporter’s chaperon. The displacement of carbonic anhydrase from the transporter has been termed “metabolon disruption” [54]. As “metabolon disruptor” we choose an antibody which binds to an epitope in the Ig1 domain of CD147, which is situated close to the CAIX binding site at position 73 (Figure 8 A). Preincubation with 1 μg/ml Anti-CD147 fully abolished the CAIX-mediated increase in MCT4 transport activity of MCT4+rCD147+CAIX-coexpressing *Xenopus* oocytes (Figure 8 B, C). Binding of Anti-CD147 to CD147 did neither influence MCT4 transport activity in the absence of CAIX (Figure 7 C) nor decrease CAIX catalytic activity (Figure 8 D). Furthermore, incubation of oocytes with Anti-CD147 induced no changes in the expression levels of MCT4, CAIX, and CD147 (Figure 9). Interestingly, Anti-CD147 recognized CD147 also in native oocytes (Figure 9 E, F). Since *Xenopus* oocytes endogenously express an analogue of CD147 [55] it appears plausible that the signal for CD147 in native oocytes derived from endogenous *Xenopus* CD147.

**Figure 8:**
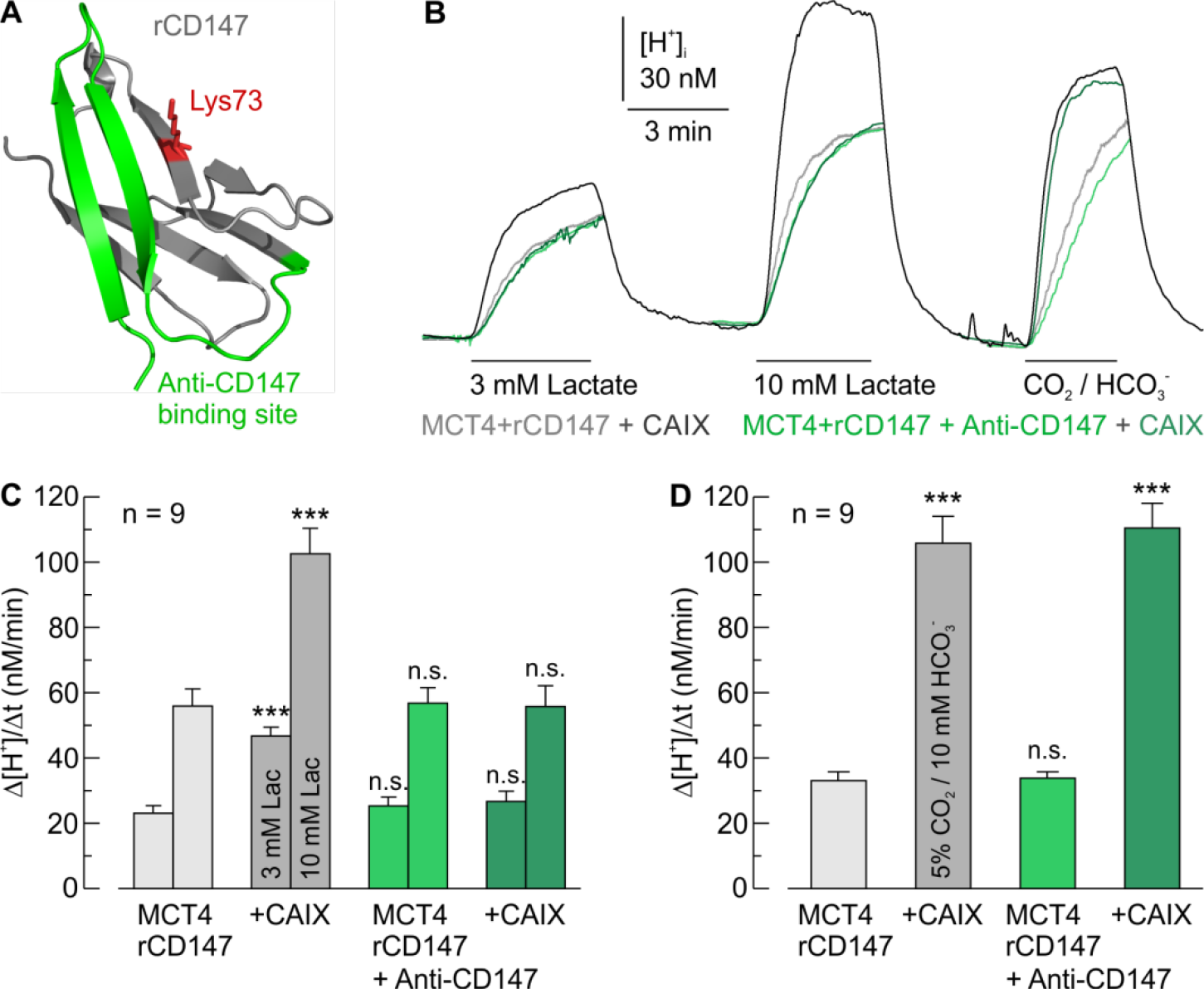
CAIX-mediated facilitation of MCT4 transport activity is disrupted by an antibody against the Ig1 domain of CD147. (**A**) Structure of the Ig1 domain of rat CD147 (PDB-ID: 3B5H [89]). The CAIX binding site K73 is labelled in red. The antibody binding region is labelled in green. (**B**) Original recordings of the change in intracellular H^+^ concentration during application of lactate and CO_2_/HCO_3_^−^ in rMCT4+rCD147-WT-coexpressing *Xenopus* oocytes, pre-incubated without (gray traces) or with an antibody against CD147 (Anti-CD147; green traces), either without (light traces) or with CAIX (dark traces). (**C**, **D**) Rate of change in intracellular H^+^ concentration (Δ[H^+^]/Δt) during application of lactate (**C**) and 5% CO_2_ / 10 mM HCO_3_^−^ (**D**), respectively, in rMCT4+rCD147-WT-coexpressing *Xenopus* oocytes, pre-incubated without (gray bars) or with Anti-CD147 (green bars), either without (light bars) or with CAIX (dark bars). The significance indicators refer to the bars for MCT4+rCD147-expressing oocytes.

**Figure 9:**
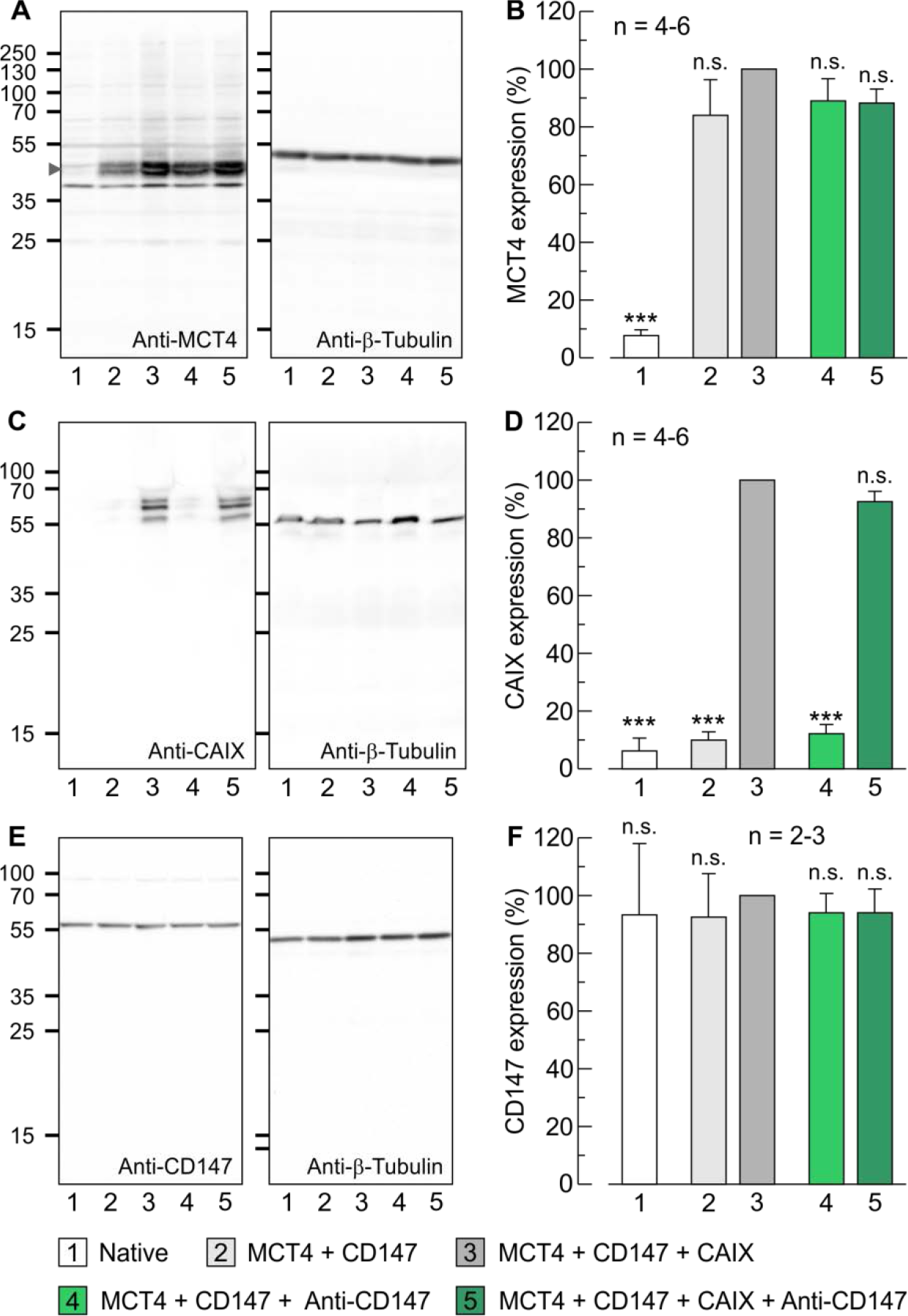
Control of protein expression in MCT4+rCD147±CAIX-expressing oocytes, pre-incubated with or without Anti-CD147. (**A**, **C**, **E**) Representative western blots against MCT4 (**A**), CAIX (**B**), and CD147 (**C**) and corresponding staining against β-tubulin (left blots) from oocytes expressing MCT4+CD147±CAIX, pre-incubated with Anti-CD147 or not, and native oocytes as control. (**B**, **D**, **F**) Relative fluorescent signal of MCT4 (**B**), CAIX (**D**), and CD147 (**F**). For every lane, the signals were normalized to the corresponding signals for β-tubulin. For every blot the signals were normalized to the signal for MCT4+CD147+CAIX-expressing oocytes.

In addition to the experiments in *Xenopus* oocytes, incubation with 10 μg/ml of Anti-CD147 resulted in a significant decrease in lactate transport capacity in hypoxic MDA-MB-231 and MCF-7 breast cancer cells, as measured by the rate of change in intracellular lactate concentration, using a lactate-sensitive FRET nanosensor (Figure 10). In MDA-MB-231 cells, Anti-CD147 decreased lactate flux by around 60% (Figure 10 C). In MCF-7 cells lactate flux was decreased by almost 65% (Figure 10 F). In both cases, disruption of the MCT-CAIX metabolon did not fully abolish MCT transport activity, since direct inhibition of MCT1/4 transport activity in MDA-MB-231 wells with α-cyano-4-hydroxycinnamic acid (CHC) and MCT1 transport activity in MCF-7 cells with AR-C155858 (AR-C) resulted in a significant reduction in lactate flux, as compared to preincubation with Anti-CD147 (Figure 10 C, F).

**Figure 10:**
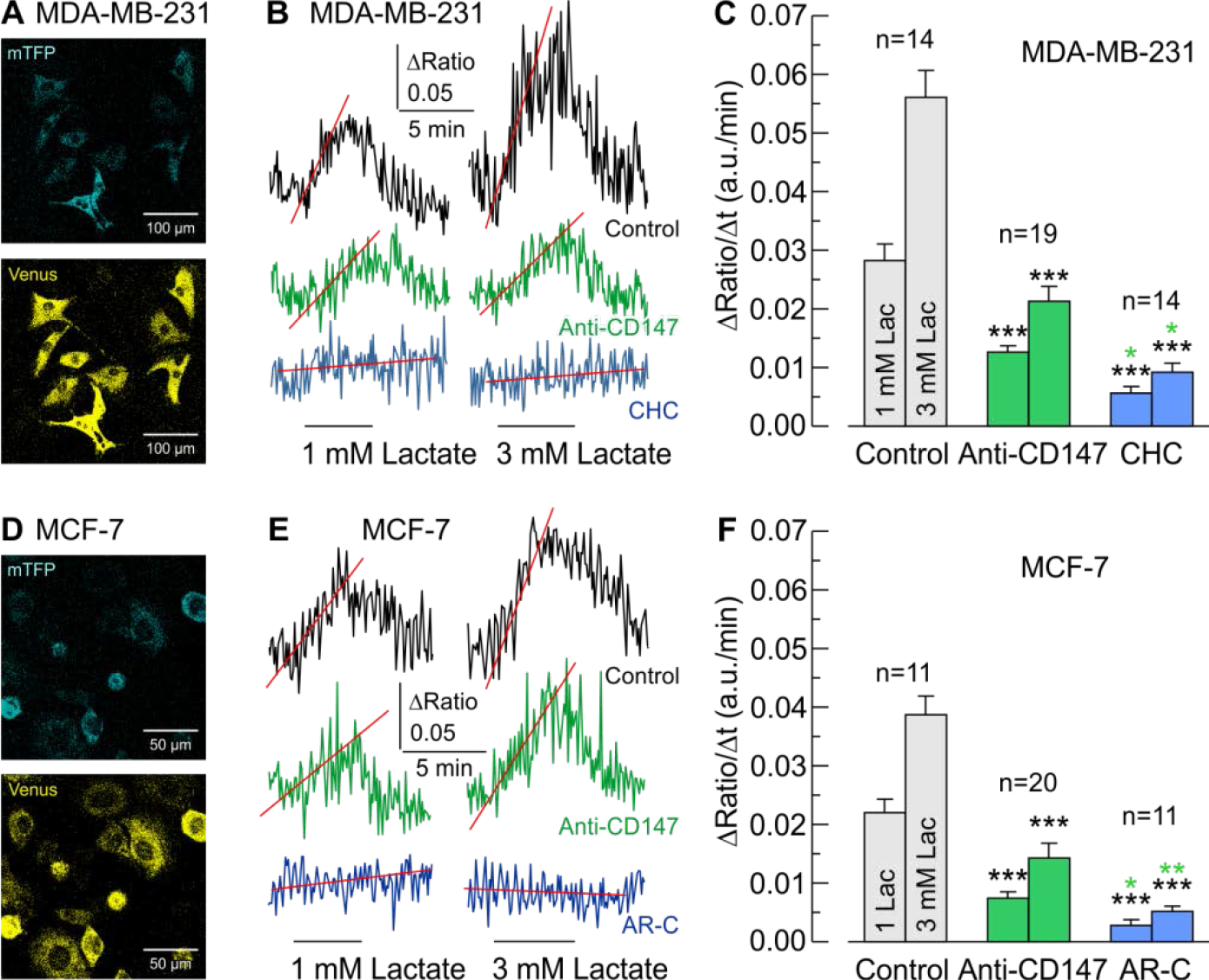
Anti-CD147 reduces lactate transport capacity in hypoxic breast cancer cells. (**A**, **D**) Fluorescent signals for mTFP (460-500 nm) (upper panel, blue) and Venus (520-550 nm) (lower panel, yellow) from MDA-MB-231 (**A**) and MCF-7 cells (**D**), transfected with the lactate-sensitive FRET nanosensor *Laconic*. (**B**, **E**) Original recordings of the relative change in intracellular lactate concentration during application of 1 and 3 mM lactate in hypoxic MDA-MB-231 (**B**) and MCF-7 cells (**E**), pre-incubated without (black and blue traces) or with Anti-CD147 (green traces), in the absence (black and green traces) or presence of the MCT inhibitors α-cyano-4-hydroxycinnamate (CHC) or AR-C155858 (blue traces). (**C**, **F**) Rate of change in lactate concentration (ΔRatio/Δt) during application of lactate in hypoxic MDA-MB-231 (**C**) and MCF-7 cells (**F**), pre-incubated without (black and blue bars) or with Anti-CD147 (green bars), in the absence (black and green bars) or presence of CHC or AR-C (blue bars). The asterisks of a given color refer to the bars of the corresponding color.

To investigate the impact of the Anti-CD147-induced metabolon disruption on cancer cell metabolism, we measured acid production in hypoxic MDA-MB-231 and MCF-7 cells in the presence of Anti-CD147 or the MCT inhibitors CHC and AR-C by determining the change in extracellular pH using a multi-well cell culture plate with integrated pH sensor (Figure 11 A-D). Application of Anti-CD147 decreased the rate of extracellular acidification by 40% in MDA-MB-231 cells and 30% in MCF-7 cells, while full inhibition of MCT transport activity with CHC (application of which resulted in a substantial acidification of the cell culture medium) or AR-C resulted in a reduction of acidification by around 80% (Figure 11 A-D). In line with these results, Anti-CD147 reduced lactate production, as determined by measuring lactate concentration in the culture medium after 24 h, by 48% in MDA-MB-231 and 38% in MCF-7 cells (Figure 11 E, F). Full inhibition of MCT transport activity with CHC and AR-C, reduced lactate production by 76% and 53%, respectively (Figure 11 E, F).

**Figure 11:**
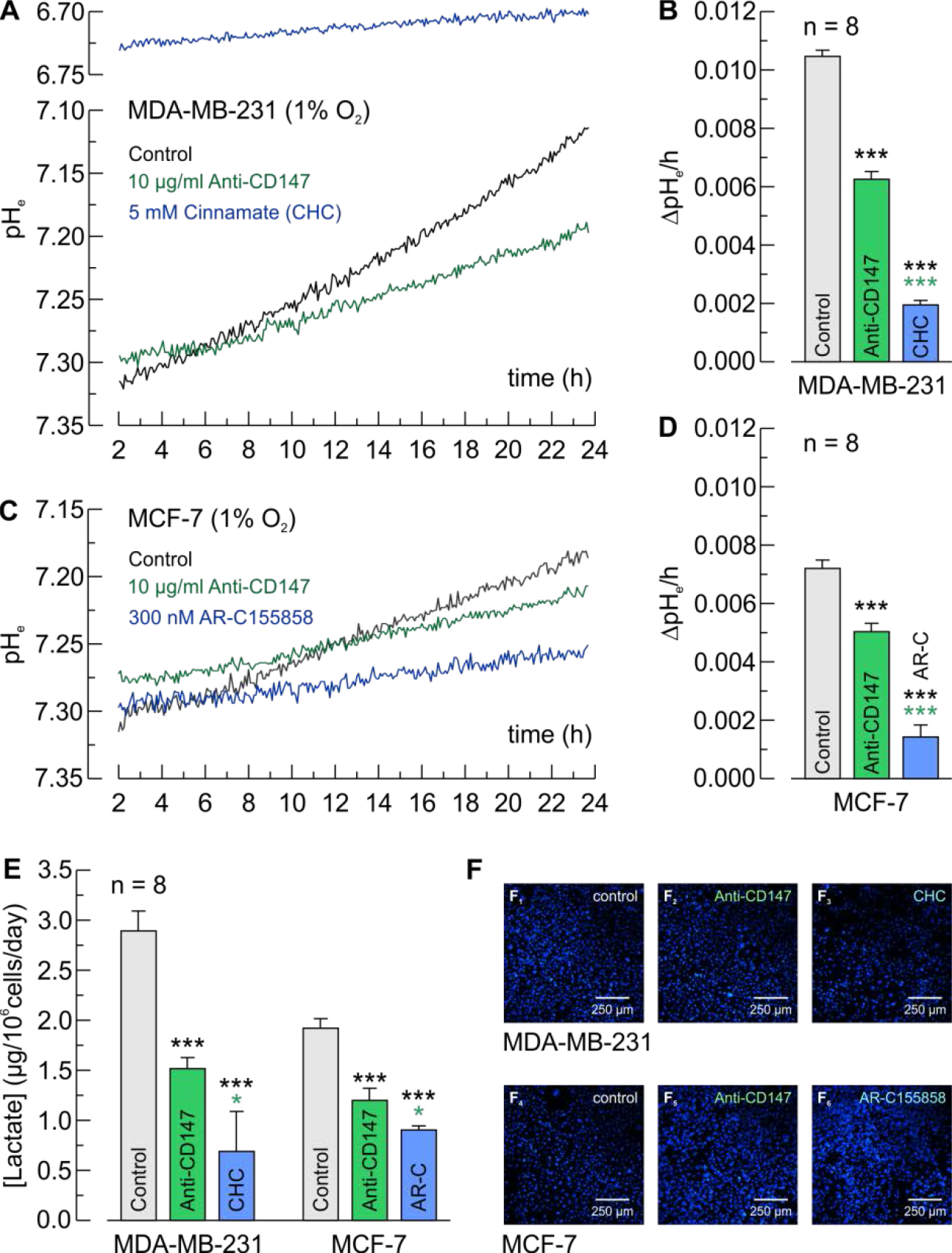
Anti-CD147 reduces glycolytic activity in hypoxic breast cancer cells. (**A**, **C**) Original recordings of extracellular pH, as measured with a SensorDish Reader in the culture medium, in hypoxic MDA-MB-231 (**A**) and MCF-7 cells (**C**), pre-incubated without (black and blue traces) or with Anti-CD147 (green traces), in the absence (black and green traces) or presence of the MCT inhibitors α-cyano-4-hydroxycinnamate (CHC) or AR-C155858 (blue traces). (**B**, **D**) Rate of change in extracellular pH (ΔpH/h) in hypoxic MDA-MB-231 (**B**) and MCF-7 cells (**D**), pre-incubated without (black and blue bars) or with Anti-CD147 (green bars), in the absence (black and green bars) or presence of CHC or AR-C (blue bars). The asterisks of a given color refer to the bars of the corresponding color. (**E**) Lactate production (in μg of lactate/10^6^cells/day), as measured with an enzymatic assay in the culture medium of hypoxic MDA-MB-231 and MCF-7 cells, pre-incubated without (black and blue bars) or with Anti-CD147 (green bars), in the absence (black and green bars) or presence of CHC or AR-C (blue bars). The asterisks of a given color refer to the bars of the corresponding color. (**F**) Representative pictures of nuclei staining of the cells used for the lactate assay, shown in (E). For every batch of cells lactate concentration was normalized to the amount of cells.

To investigate whether metabolon disruption with Anti-CD147 has an impact on cancer cell proliferation we determined the number of hypoxic MDA-MB-231 and MCF-7 cells for up to three days in the presence of 10 μg/ml of Anti-CD147 or CHC and AR-C (Figure 12). Anti-CD147 reduced proliferation of hypoxic MDA-MB-231 and MCF-7 cells by 62%-94% and 67%-85%, respectively (Figure 12 C, D). Inhibition of MCT1 transport activity with AR-C decreased proliferation of MCF-7 cells by 82%-88%, while CHC almost fully abolished proliferation of MDA-MB-231 cells (Figure 12 C, D). Application of Anti-CD147 at a concentration of only 5 μg/ml resulted in a lower reduction in proliferation of MDA-MB-231 (22%-50%) and MCF-7 cells (45%-68%) (Figure 13).

**Figure 12:**
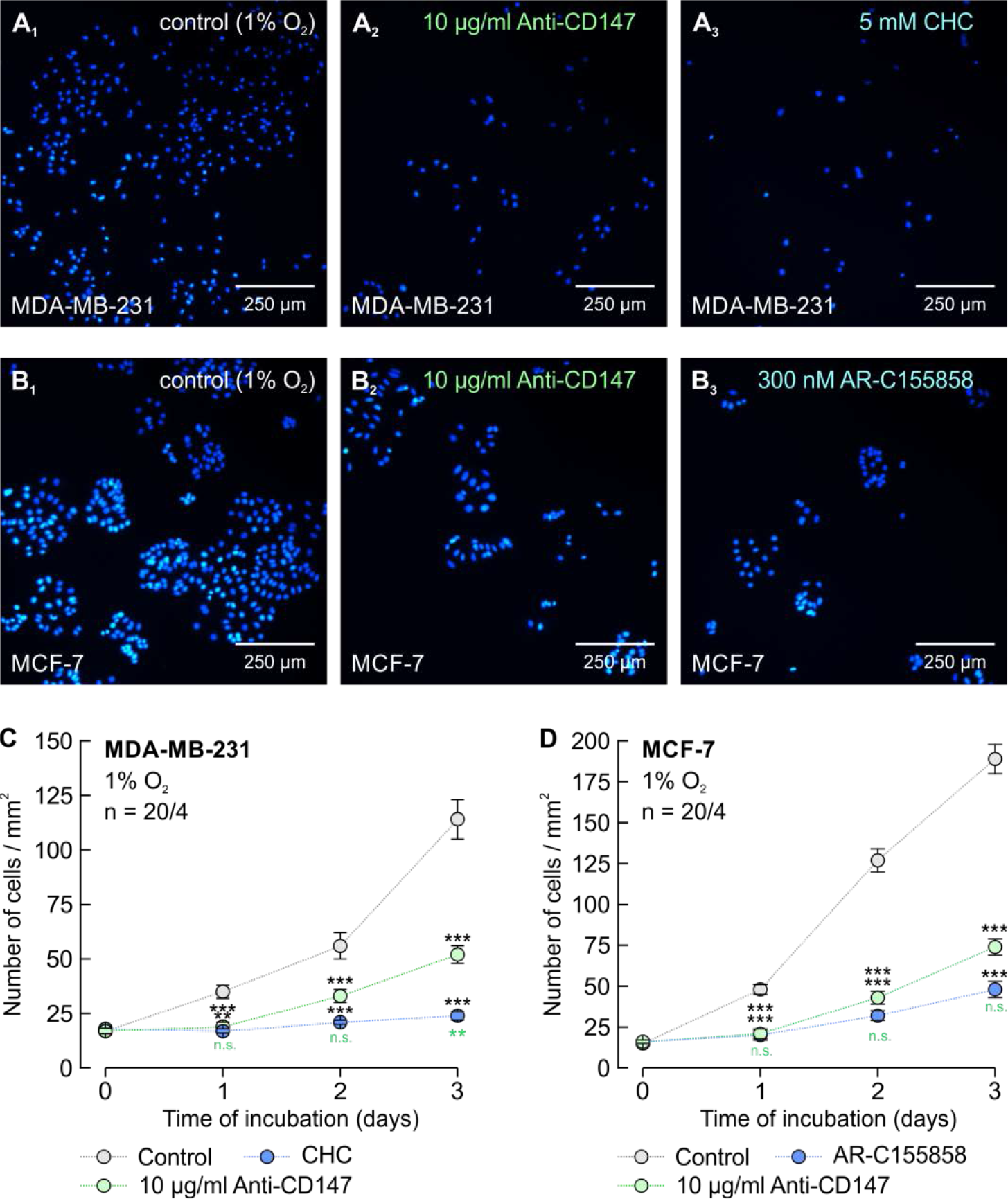
Application of 10 μg/ml of Anti-CD147 reduces proliferation of hypoxic breast cancer cells. (**A, B**) Staining of nuclei with Hoechst 33342 (blue) in MDA-MB-231 (**A**) MCF-7 cells (**B**), after 3 days in culture. Hypoxic cells remained either untreated (**A**_**1**_, **B**_**1**_), incubated with 10 μg/ml Anti-CD147 (**A**_**2**_, **B**_**2**_), or incubated with the MCT inhibitors α-cyano-4-hydroxycinnamate (CHC) or AR-C155858 (**A**_**3**_, **B**_**3**_). (**C**, **D**) Total number of nuclei/mm^2^ in MDA-MB-231 (**C**) and MCF-7 cell cultures (**D**), kept for 0–3 days under the conditions described in (A, B). For every data point four dishes of cells were used and five pictures were taken from each dish at random locations, yielding 20 pictures/data point (n = 20/4).

**Figure 13:**
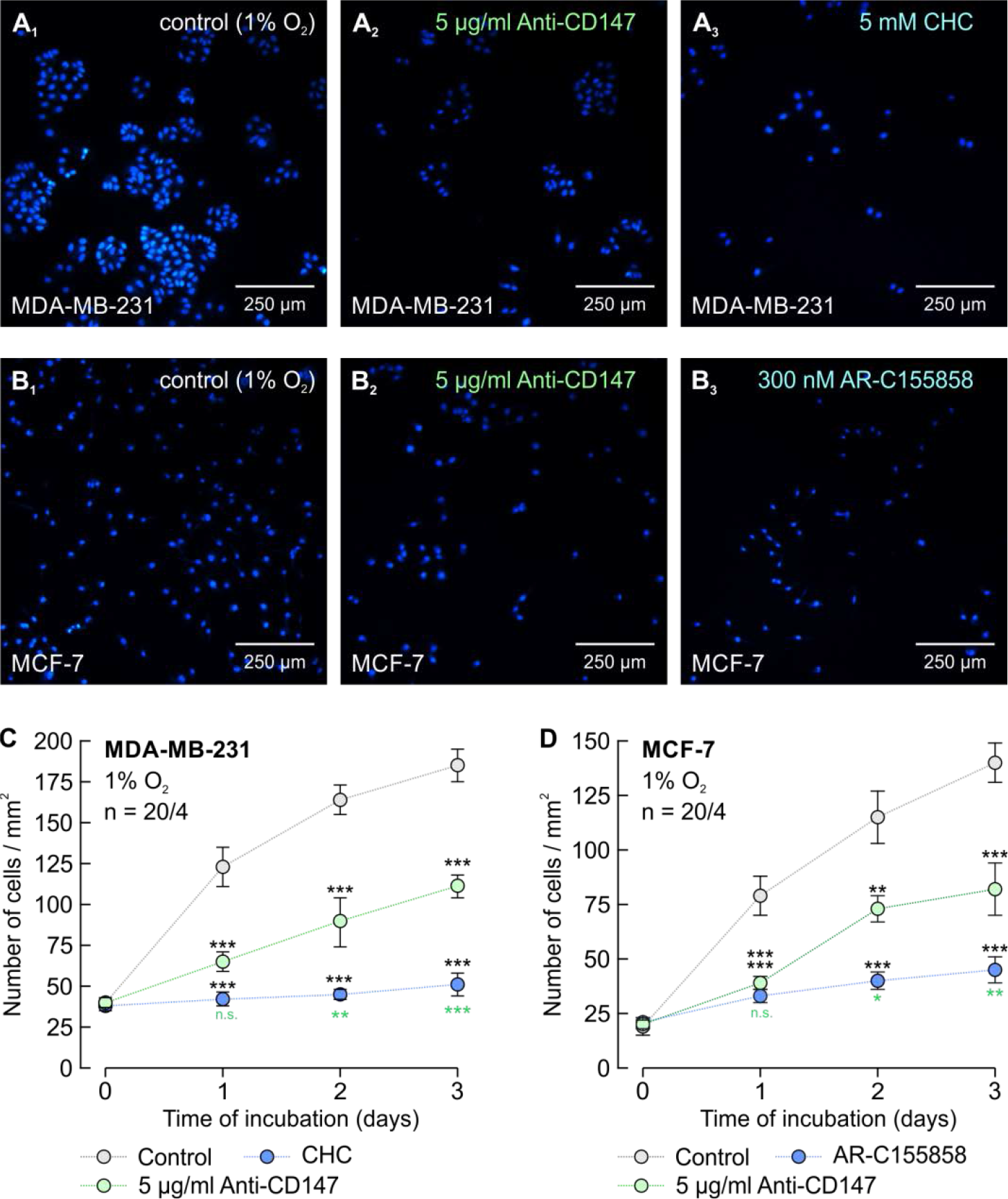
Application of 5 μg/ml of Anti-CD147 reduces proliferation of hypoxic breast cancer cells. (**A, B**) Staining of nuclei with Hoechst 33342 (blue) in MDA-MB-231 (**A**) MCF-7 cells (**B**), after 3 days in culture. Hypoxic cells remained either untreated (**A**_**1**_, **B**_**1**_), incubated with 5 μg/ml Anti-CD147 (**A**_**2**_, **B**_**2**_), or incubated with the MCT1 inhibitors α-cyano-4-hydroxycinnamate (CHC) or AR-C155858 (**A**_**3**_, **B**_**3**_). (**C**, **D**) Total number of nuclei/mm^2^ in MDA-MB-231 (**C**) and MCF-7 cell cultures (**D**), kept for 0–3 days under the conditions described in (A, B). For every data point four dishes of cells were used and five pictures were taken from each dish at random locations, yielding 20 pictures/data point (n = 20/4).

Taken together, these results demonstrate that disruption of the MCT1/4-CD147-CAIX transport metabolon with an antibody against the Ig1 domain of CD147 can significantly decrease lactate transport over the cell membrane, which results in a decrease of lactate and proton production (synonymic with a decrease in glycolytic activity) and therefore a decrease in cancer cell proliferation.

## Discussion

The present study shows for the first time that carbonic anhydrase IX forms a transport metabolon with the two major H^+^/lactate extruders MCT1 and MCT4 in human tumor tissue, but not in healthy breast tissue. The number of transport metabolons increased with higher tumor grade, with the number of MCT1-CAIX metabolons increasing from grade I to grade II and the number of MCT4-CAIX metabolons increasing from grade II to grade III. In line with this, others have shown that CAIX is expressed in breast tumors, but absent in healthy breast tissue [56]. Furthermore, expression of CAIX, MCT1, and MCT4 have all been shown to positively correlate with tumor grade and poor prognosis [2, 10, 12–14, 34–40].

It was recently demonstrated by sequential PET/CT and MRI that glycolytic activity increases with higher tumor grade in breast cancer patients [57]. These results are in agreeance with our findings that the number of MCT1/4-CAIX transport metabolons increases with higher tumor grade in breast tumor tissue samples. Since a higher glycolytic rate results in increased production of lactate and protons, these cells would also require a higher H^+^/lactate efflux capacity, which could be met by an increased amount of MCT1/4-CAIX transport metabolons in the cell membrane. Interestingly, the ratio between MCT1-CAIX and MCT4-CAIX metabolons seems to change with higher tumor grade. MCT4 displays a lower affinity for lactate than MCT1, but has a higher transport capacity [6–8]. Therefore MCT4, the expression of which was found to be upregulated in cancer cells under hypoxia [58], is considered a high capacity lactate exporter, while MCT1 was suggested to serve both as a lactate importer and exporter in cancer cells, depending on the cell’s metabolic profile [59, 60]. It could be speculated that in low grade tumors, which show lower overall glycolytic activity, a glycolytic population of cells, which is situated in the hypoxic regions of the tumor, might function as lactate exporters, while another population of cells takes up lactate for oxidative energy production, as previously hypothesized [59]. In that case moderate lactate export and import could be mediated by MCT1 in cooperation with CAIX. With increasing glycolytic activity, as observed with higher tumor grade, cells would require maximum efflux capacity for lactate and protons, which could only be met by a transport metabolon formed by MCT4 and CAIX.

The necessity for an increasing amount of MCT1/4-CAIX transport metabolons in higher grade tumors could also derive from progressing restriction of acid removal. MRI studies on breast cancer patients demonstrated that the apparent diffusion coefficient (ADC) decreased with higher tumor grade [57, 61, 62]. It can be assumed that a lower diffusion coefficient indicates reduced venting of lactate and protons from the tumor mass. This reduction would result in accumulation of both ions in tumor extracellular space, which, in turn, would create an unfavorable gradient for proton-coupled lactate export. Especially diffusive flux of the highly-buffered protons could be significantly restricted in tumor tissue [63]. In such an environment, efficient proton handling seems crucial for proton-coupled lactate transport across the membrane. We have previously suggested that intracellular and extracellular carbonic anhydrases can function as “proton antennae” for MCTs by mediating the rapid exchange of protons between the transporter pore and the surrounding protonatable residues [29, 64–67]. In CAIX, proton transfer between MCT and surrounding protonatable residues seems to be mediated by acidic residues within the enzyme’s proteoglycan-like domain [30]. Rapid H^+^ transfer between protonatable sites can be mediated by electrostatic repulsion caused by overlapping Coulomb cages, which would require a maximum distance below 1 nm between the involved residues [68–70]. Such a close proximity could be achieved by direct binding of the proteins. Previous studies have shown that intracellular CAII binds to a cluster of three glutamic acid residues in the C-terminal tail of MCT1 (E^489^EE) and MCT4 (E^431^EE), respectively [71, 72]. Extracellular CAs, however, do not directly interact with the transporter but indirectly via the transporter’s chaperons. We recently showed that CAIV, the catalytic domain of which is tethered to the extracellular face of the plasma membrane via a glycosylphosphatidylinositol (GPI) anchor, binds to Glu73 in the Ig1 domain of CD147 [49]. CAIV binding is mediated by His88, which also serves as central residue of the enzyme’s intramolecular proton shuttle [49, 73]. In the present study, we demonstrate that CAIX binds to the Ig1 domain of CD147 (and GP70). CD147 binding is mediated by Glu73 (Lys73 in rat CD147 and Arg130 in rat/human GP70), while in CAIX binding requires His200. Molecular docking shows that CD147-Glu73 can form a hydrogen bond with CAIX-His200 in the “in” confirmation, with a distance of 2.1 Å between the two binding partners (Figure 14 A). Binding of the chaperon to CAIX-His200 is mediated either by an acidic residue (hCD147-Glu73), which serves as proton acceptor in a hydrogen bond, or an alkaline residue (rCD147-Lys73 or r/hGP70-Arg130), which serves as proton acceptor. Therefore CAIX-His200 must either serve as proton donor or proton acceptor, depending on its binding partner. We could previously show that CAIV-His88 (the homologue to CAIX-His200) which binds to the Ig1 domain of CD147 and GP70, can indeed either serve as hydrogen donor or hydrogen acceptor, depending on whether it binds to hCD147-Glu73 or to rCD147-Lys73 / hGP70-Arg130 [49]. Therefore, it can be assumed that CAIX-His200 also forms a hydrogen bond with hCD147-Glu73 or rCD147-Lys73 / hGP70-Arg130. This binding would bring CAIX close enough to the transport pore to establish an efficient proton shuttle between transporter and enzyme. Indeed, mutation of either rCD147-Lys73 or CAIX-His200 does not only result in a loss of binding between CD147 and CAIX, but also in a loss of functional interaction between MCT1/4 and CAIX (for CAIX-His200, see Figure 4 in [29]). Interestingly, CAIX-His200, which serves as binding site for CD147/GP70, does also represent the central residue of the enzyme’s intramolecular proton shuttle. We recently showed for intracellular CAII, that CAII-His64 (the central residue of the enzyme’s intramolecular H^+^ shuttle), mediates binding of the enzyme to an acidic cluster in the MCT1/4 C-terminal tail, but is not involved in the transfer of protons between transporter and enzyme [66]. This proton transfer is instead mediated by the two acidic residues CAII-Glu69 and CAII-Asp72 at the protein surface [66]. The catalytic domain of CAIX does not seem to feature a homologue cluster to CAII-Glu69 and CAII-Asp72, however, the proteoglycan-like domain of CAIX features 18 glutamate and 8 aspartate residues, which have been suggested to function as an intramolecular proton buffer [74]. Truncation of the PG domain resulted in a loss of functional interaction between MCT1/4 and CAIX [30] even though in this study, binding of CAIX to CD147 persisted after truncation. Taken together, the data indicate that CAIX, which binds to the Ig1 domain of the MCT chaperon CD147 (or GP70) via CAIX-His200, facilitates the acidic residues in its PG domain to rapidly exchange protons between transporter pore and the surrounding protonatable residues (Figure 14 B_1_). This rapid H^+^-exchange would counteract the formation of proton-microdomains (local accumulation or depletion of protons) at the extracellular site of the transporter pore and could thereby drive proton-coupled lactate flux across the cell membrane.

**Figure 14:**
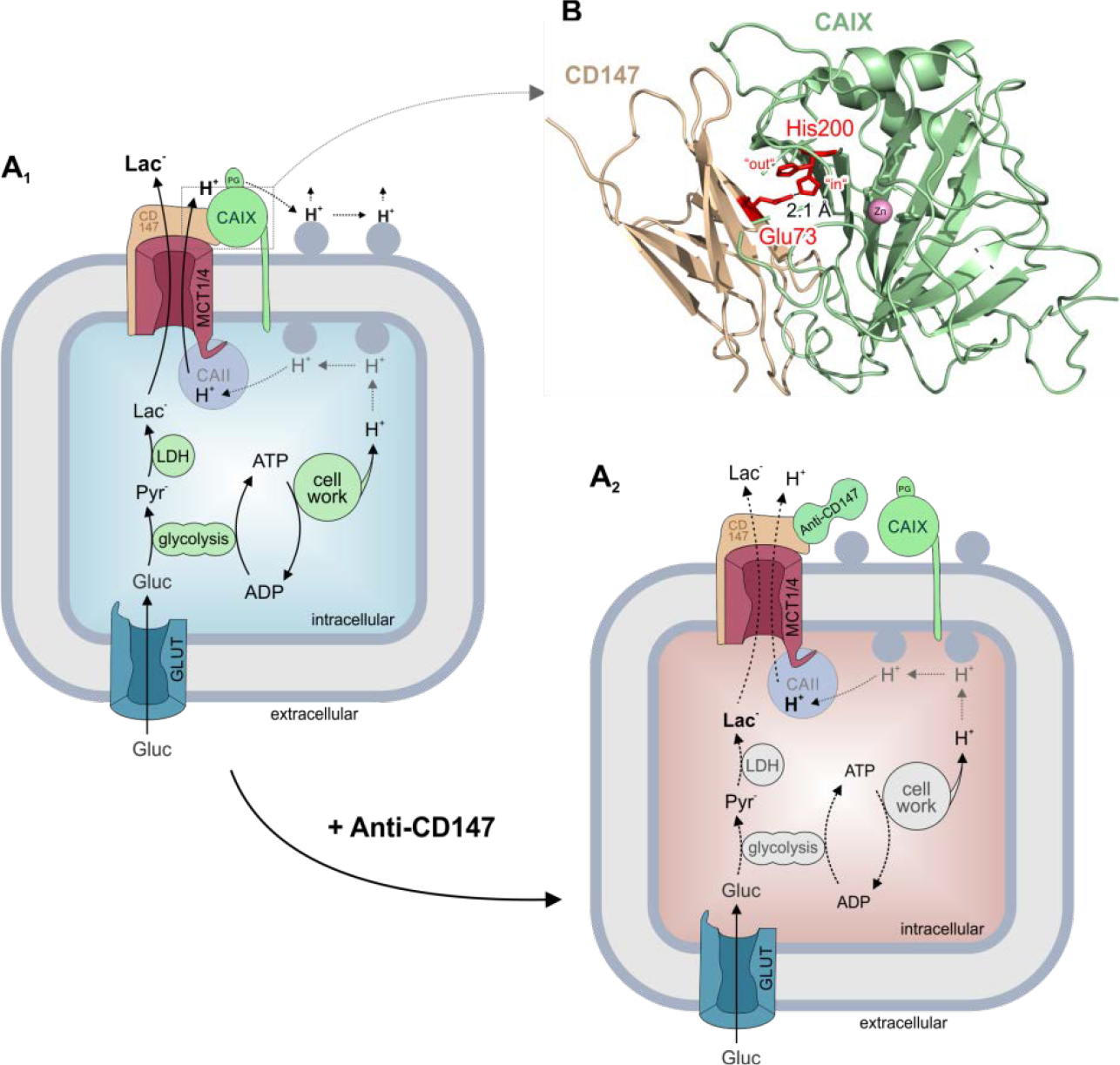
Model of the MCT1/4-CD147-CAIX transport metabolon. (**A**_**1**_) Glycolysis serves as the prime energy source for hypoxic cancer cells, leading to vast production of lactate and protons which have to be removed from the cell to avoid intracellular acidosis and suffocation of cell metabolism. Under these conditions CAIX, which is directly bound to the Ig1 domain of the MCT1/4 chaperon CD147, serves as a proton antenna for the transporter which rapidly exchanges H^+^ between transporter pore and surrounding protonatable residues (blue-gray circles) at the extracellular site of the plasma membrane. Fast removal of H^+^ counteracts the formation of proton microdomains around the transporter pore and drives the efflux of lactate and protons from the cell. Cytosolic CAII, which is directly bound to the MCT1/4 C-terminal tail, serves as the intracellular counterpart to CAIX. CAII collects H^+^ from surrounding protonatable residues at the inner face of the plasma membrane and shuttles them to the transporter pore. (**A**_**2**_) Binding of Anti-CD147 to an epitope close to the CAIX binding site in the Ig1 domain of CD147, drives CAIX away from the MCT1/4-CD147 complex. This disruption of the transport metabolon leads in a decrease in MCT1/4 transport capacity, resulting in intracellular accumulation of lactate and protons, which in turn leads to a decrease in glycolytic activity and ultimately a reduction in cell proliferation. (**B**) Structural model of the physical interaction between CAIX and CD147. CAIX (green structure) binds to the Ig1 domain of CD147 (ochre structure) by formation of a hydrogen bond (dotted line) between CD147-Glu73 and CAIX-His200 in the “in” confirmation (red sticks), with a distance of 2.1 Å between the two binding partners.

Inhibition of CAIX catalytic activity has been widely considered as a potential therapeutic strategy for hypoxic tumors [75]. However, most of the inhibitors tested so far bind to a moiety apart from the His200 and might therefore not target the direct and functional interaction between MCT1/4-CD147 and CAIX [76]. To test whether disruption of the MCT-CD147-CAIX transport metabolon could impact on cancer cell metabolism and decrease cell proliferation we used an antibody against CD147 (Anti-CD147), which targets a moiety close to the CD147-Glu73. Application of Anti-CD147 resulted in the total loss of CAIX-induced increase in MCT4 transport activity in *Xenopus* oocytes, indicating that the antibody does indeed disrupt the interaction between MCT1-CD147 and CAIX. In MCF-7 and MDA-MB-231 cancer cells, disruption of the transport metabolon resulted in a significant rate reduction, but no full inhibition of lactate flux. This is not surprising as MCT transport activity is increased by carbonic anhydrases, but MCTs are already active in the absence of the enzyme. However, disruption of the transport metabolon with Anti-CD147 resulted in a significant decrease in lactate and proton production and ultimately a decrease in cell proliferation. These results indicate that disruption of the transport metabolon results in a decrease of MCT transport activity, which leads to intracellular lactate accumulation and inhibition of glycolytic activity, which in turn results in decreased cell proliferation (Figure 14 B_2_).

Direct inhibition of MCT transport activity by small molecule inhibitors like AR-C155858 or AZD3965 is considered a therapeutic strategy for tumor treatment [52, 53, 77]. Since metabolon disruption only reduces, not fully abolishes MCT transport activity, direct inhibition of MCT1 and MCT4 seems more efficient for cancer treatment on the cellular level. However, MCTs are ubiquitously expressed in the human body and play a central function in the energy metabolism of a wide range of tissues, including heart and skeletal muscle, liver and brain [77–80]. Therefore, systemic application of a high dose of MCT inhibitors, which would be required for full inhibition of MCT transport activity in a tumor, could be expected to produce severe side effects in other tissues. Therefore, systemic administration of a MCT inhibitor could also only aim on reduction, but not full inhibition of MCT transport activity within the tumor. CAIX, however, is expressed in few healthy tissues, including stomach and gallbladder, but highly upregulated in many solid tumors [32]. Therefore, targeted disruption of the MCT-CD147-CAIX transport metabolon should cause fewer side effects in other tissue than direct targeting of MCT transport function. However, such targeted inhibition has to be directed against CAIX, which is the only part of the MCT-CD147-CAIX metabolon that is exclusively expressed in cancer cells. We therefore suggest a screen for small molecule inhibitors or antibodies against CAIX, which could interfere with binding of the enzyme to the Ig1 domain of CD147, to selectively reduce lactate transport capacity in cancer cells, without interfering with lactate flux in healthy tissue.

Taken together the results provide a proof of concept that the MCT1/4-CD147-CAIX transport metabolon, found in breast cancer tissue, can be exploited as a potential drug target to interfere with cancer cell’s energy metabolism in order to reduce cell proliferation and thereby tumor progression.

## Materials and Methods

### Tissue micro array

Human breast tissue micro arrays were purchased from Novus Biological (NBP2-47174). The microarrays contain 16 cases of breast cancer, each in duplicates, with corresponding uninvolved tissue from the same patient as control (48 cores, 2 mm in diameter, 4 μm thick). According to the manufacturer, all tissues were from surgical resection, fixed in 10% neutral buffered formalin for 24 hours and processed using identical standard operating procedures. The specifications of the tumor samples are given in Table 1.

**Table 1:**
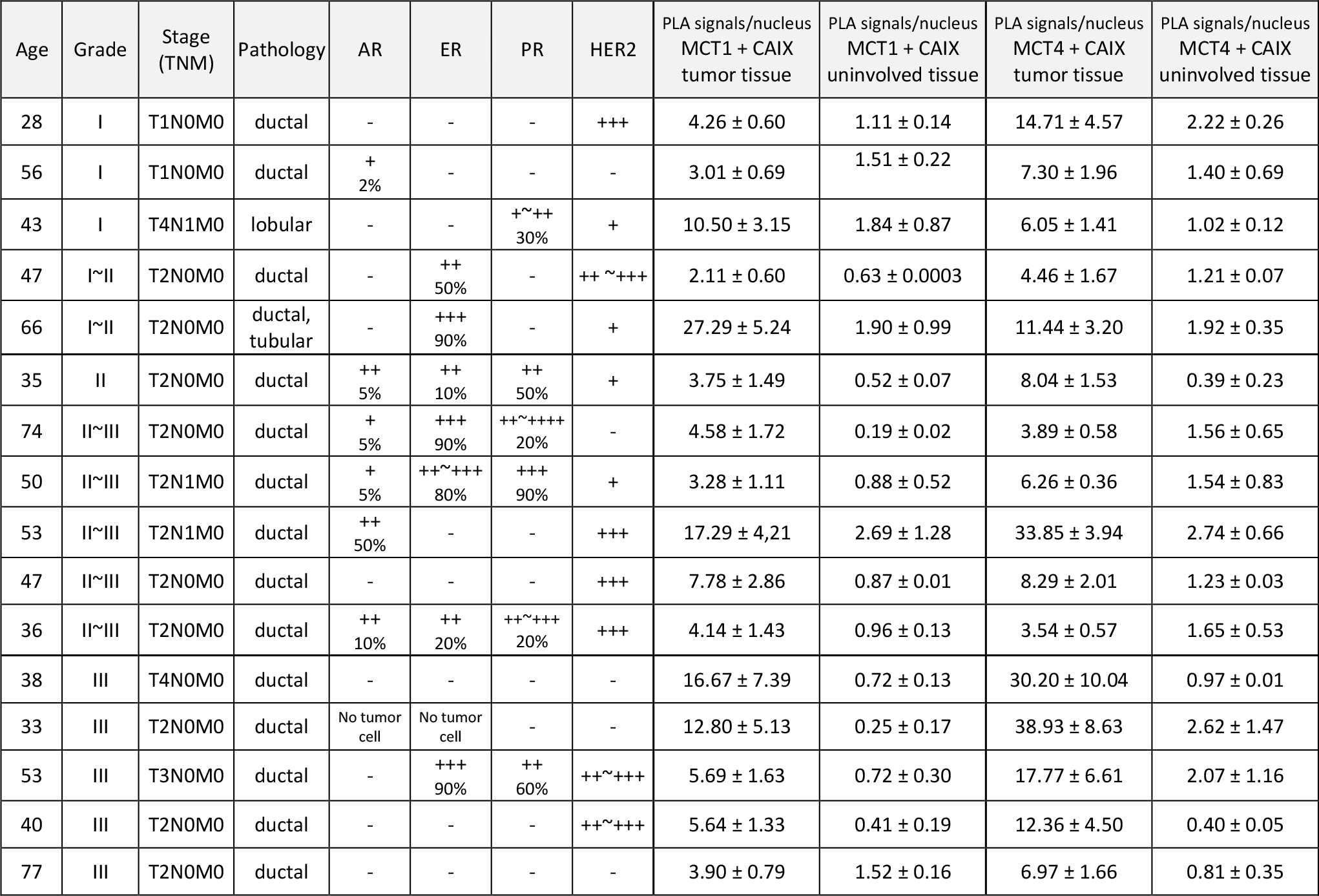
Specifications of the tissue microarray and number of PLA signals. The tissue micro array was purchased form Novus Biological (Human breast tissue microarray (Tumor), NBP2-47174). The array contains 16 cases of breast cancer (each in duplicates) paired with one sample of adjacent normal tissue. All patients were female. Stage classification: T: Primary Tumor; “T1” Tumor 2 cm or less in greatest dimension; “T2” Tumor more than 2 cm but not more than 5 cm in greatest dimension; “T3” Tumor more than 5 cm in greatest dimension; “T4” Tumor of any size with direct extension to chest wall or skin. N: Regional Lymph Nodes; “N0” No regional lymph node metastasis; “N1” Metastasis to movable ipsilateral lymph node(s). M: Distant Metastasis; “M0” No distant metastasis. Pathology: “ductal” invasive ductal carcinoma; “tubular” invasive lobular carcinoma; “ductal, lobular” invasive ductal carcinoma, tubular type. Receptor type: “AR” androgen receptor; “ER” estrogen receptor; “PR” progesterone receptor; “HER2” human epidermal growth factor receptor 2 (receptor tyrosine-protein kinase erbB-2, CD340). Staining scoring: “−” no staining; “+” weak staining; “++” moderate staining; “+++” strong staining; “%” percentage of positive cells. The PLA signals/nucleus derive from two pictures, taken at random locations in each sample. Tumor samples were present in duplicates with one sample of adjacent normal tissue.

### Cultivation of breast cancer cell lines

The human breast adenocarcinoma cell lines MCF-7 (DSMZ-No. ACC-115) and MDA-MB-231 (DSMZ-No. ACC-732) were purchased from the German Collection of Microorganisms and Cell Cultures (DSMZ, Braunschweig, Germany). If not mentioned otherwise, both cell lines were cultured in RPMI-1640 medium with L-glutamine and sodium bicarbonate (R8758, Sigma-Aldrich, Germany), supplemented with 10% fetal bovine serum (F7524, Sigma-Aldrich), and 1% penicillin-streptomycin (P4333, Sigma-Aldrich). Cells were either incubated under normoxia (95% air, 5% CO_2_) or under hypoxia (94% N_2_, 5% CO_2_, 1% O_2_) in humidified cell culture incubators. Both cell lines were subcultivated for a maximum of 15 passages and tested negative for contamination with mycoplasma.

### Interfering antibodies and inhibitors

Mouse monoclonal Anti-CD147 IgG_1_, directed against amino acids 13-51 within the Ig1 domain of human CD147 was purchased from Santa Cruz Biotechnology (sc-374101). Before the antibody was used in cell culture or on *Xenopus* oocytes, NaN_3_ was removed by exchanging the buffer solution for sterile PBS, using a protein concentrator centrifuge column with a molecular cut-off of 10 kDa (Pierce™ Protein Concentrator PES, 10K MWCO, 2-6 ml; No. 88516). *Xenopus* oocytes (kept in saline without serum) were incubated with 1 μg/ml of Anti-CD147 in oocyte saline for 24 hours. If not mentioned otherwise, MCF-7 and MDA-MB-231 cells were incubated with 10 μg/ml of Anti-CD147, to account for unspecific binding of the antibodies to compounds in the culture medium. Cells were incubated for 24 hours to ensure maximum binding of the antibodies. The MCT1 inhibitor AR-C15585 was purchased from Tocris Bioscience (No. 4960). The inhibitor was dissolved at a concentration of 10 mM in dimethyl sulfoxide (DMSO) and filtered sterile. AR-C155858 was used at a working contraction of 300 nM. The generic MCT inhibitor α-cyano-4-hydroxycinnamic acid (CHC) was purchased from Sigma-Aldrich (No. 476870). CHC was dissolved in DMSO at a concentration of 1 M and filtered sterile. The inhibitor was used at a working concentration of 5 mM.

### *In situ* proximity ligation assay

Interaction between MCT1, MCT4 and CAIX in human breast cancer tissue micro arrays, as well as in cultivated MCF-7 and MDA-MB-231 cells, was examined using the Duolink^®^ *in situ* Proximity Ligation Assay (PLA) Kit (DUO92103, Sigma-Aldrich). The protocol for the PLA in cultivated breast cancer cells has been described in detail previously (Noor et al., 2018). MCF-7 cells where cultured in RPMI medium as described above. MDA-MB-231 cells were cultured in Leibovitz-L15 medium (No. 11415; ThermoFisher), supplemented with 10% fetal calf serum (F413, Sigma-Aldrich), 1% penicillin/streptomycin (15149, Thermo Fisher), and 5 mM D-glucose (16325, Sigma-Aldrich). The PLA was carried out with the following antibodies: goat anti-MCT1 polyclonal antibody (T-19), sc-14917, Santa Cruz Biotechnology, 1 μg/ml; goat anti-MCT4 polyclonal antibody (C-17), sc-14930, Santa Cruz Biotechnology, 1 μg/ml; mouse anti-CAIX monoclonal antibody (M75), kindly provided by Dr. Silvia Pastorekova [81], 1 μg/ml. For better visualization of cells, actin filaments were stained with Alexa Fluor 488-labelled phalloidin (1:500; A12379; Life Technologies). Nuclei were stained with 4′,6-diamidino-2-phenylindole (DAPI), added to the mounting medium (DUO82040, Sigma-Aldrich). Pictures were taken with a Zeiss LSM700 confocal laser scanning microscope, using a 40x oil immersion objective (EC Plan-Neofluar 40x/1.3; Carl Zeiss AG). For each sample, seven pictures were taken at random locations.

For dewaxing of the tissue micro array prior to staining, the slides were backed for 30 min at 60°C and washed three times with xylene. The procedure was repeated until samples showed a white color without any inclusions. Afterwards, slides were rehydrated by incremental 10 min incubation with decreasing alcohol concentration (100%, 90%, 70%, 50% EtOH). The tissue was incubated in deionized H_2_O for 5 min, washed 3 times with phosphate-buffered saline (PBS) and permeabilized in 0.4% Triton X-100 (T878, Sigma-Aldrich), 1% donkey serum (D9663, Sigma-Aldrich). The PLA was then carried out the same way as described for cultured cells. Pictures were taken with a Zeiss LSM700 confocal laser scanning microscope, using a 25x oil immersion objective (LD LCI Plan-Apochromat 25x/0.8 Imm Korr DIC; Carl Zeiss AG). For each tissue sample two pictures were taken at random location. Every breast cancer sample was present in duplicate with one sample of corresponding uninvolved breast tissue. PLA signals were analyzed using the software ImageJ.

### Antibody staining in cultured cancer cells

Conventional antibody staining of cultured breast cancer cells was described in detail previously [29, 66]. The antibodies used for conventional antibody staining were identical to the antibodies used for the PLA and were used in the same concentration. Pictures were taken with a Zeiss LSM700 confocal laser scanning microscope, using a 40x oil immersion objective (EC Plan-Neofluar 40x/1.3; Carl Zeiss AG).

### Western Blot analysis

Protein quantification in *Xenopus* oocytes by western blot analysis was described in detail previously [82]. In brief, oocytes were lysed in PBS with 1% ionic detergent and protease inhibitors. Lysate was cleared by centrifugation for 15 min at 15000xg, 4°C. 25 μg of total protein were separated on a 12% SDS gel and blotted to a polyvinylidene difluoride membrane (Roti-PVDF, pore size 0.45 μm, Carl-Roth GmbH). CAIX was labeled with mouse anti-CAIX monoclonal antibody (M75; [81]; 0.4 μg/ml); MCT4 was labeled with rabbit Anti-MCT4 polyclonal antibody (AB3314P, Millipore; 4 μg/ml); CD147 was labeled with mouse Anti-CD147 monoclonal antibody (sc-374101, Santa Cruz Biotechnology; 0.1 μg/ml); β-tubulin was labeled with mouse Anti-β-tubulin monoclonal antibody (T5201, Sigma-Aldrich; 2 μg/ml). Quantification of the bands was carried out using the software ImageJ. Concentrations of CAIX, MCT4, and CD147, respectively, were normalized to the concentration of β-tubulin in the same sample.

### Pull-down of CAIX with GST fusion proteins

CAIX was pulled down with GST fusion proteins of CD147 and GP70, respectively, using the Pierce™ GST protein interaction pull-down Kit (21516, Thermo Fisher), as previously described [49, 72]. In brief, GST-fusion proteins of the Ig1 domain of human CD147, rat CD147, and rat GP70, respectively, cloned into the expression vector pGEX-2T (GE Healthcare), were expressed in *E. coli* BL21 cells and coupled to glutathione-coated agarose beads. CAIX was expressed in *Xenopus* oocytes, as described in the next chapter. For each pulldown, lysate from 30 oocytes was added to the beads. The pulldown was then analyzed using western blot. CAIX was labelled with mouse anti-CAIX monoclonal antibody (M75; [81]; 0.4 μg/ml); GST was labeled with mouse Anti-GST Tag monoclonal antibody (05-782, Merck Chemicals; 2.5 μg/ml). To account for variations in the amount of fusion protein, each signal for CAIX was normalized to the corresponding signal for GST.

### Heterologous protein expression in Xenopus oocytes

Protein expression in *Xenopus* oocytes was carried out as previously described [83, 84]. In brief, cDNA coding for human CAIX, rat MCT1, rat MCT2, rat MCT4, rat CD147-WT or a mutant of the protein, as well as rat GP70-WT or mutant, all cloned into the *Xenopus* expression vector pGEM-He-Juel, was transcribed *in vitro* using the Invitrogen™ Ambion™ mMESSAGE mMACHINE™ T7 Transcription Kit (Thermo Fisher). *Xenopus laevis* females were purchased from the Radboud University, Nijmegen, Netherlands. The procedure for surgical removal of oocytes from anaesthetized frogs was approved by the Landesuntersuchungsamt Rheinland-Pfalz, Koblenz (23 177-07/A07-2-003 §6) and the Niedersächsisches Landesamt für Verbraucherschutz und Lebensmittelsicherheit, Oldenburg (33.19-42502-05-17A113). Oocytes were singularized by collagenase treatment for 1 h at 28°C in Ca^2+^-free oocyte saline. Singularized oocytes were stored in Ca^2+^-containing oocyte saline (82.5 mM NaCl, 2.5 mM KCl, 1 mM CaCl_2_, 1mM MgCl_2_, 1mM Na_2_HPO_4_, 5mM HEPES, pH 7.8) at 18°C. Oocytes of the developmental stages IV and V were injected with 5 ng of cRNA coding for MCT1, MCT2, or MCT4, together with 10 ng of cRNA coding for rCD147-WT or a mutant of rCD147, or rGP70-WT or a mutant of rGP70, and 5 ng of cRNA coding for CAIX-WT.

### Measurements of intracellular H^+^ concentrations in Xenopus oocytes

Determination of intracellular H^+^ concentrations in *Xenopus* oocytes with ion-sensitive microelectrodes was described in detail previously [84, 85]. Oocytes were clamped to a holding potential of −40 mV as previously described [84, 85]. All measurements were carried out in oocyte saline, pH 7.0, in the nominal absence of CO_2_/HCO_3_^−^, at room temperature. In lactate-containing solution, NaCl was replaced by Na-L-lactate in equimolar amounts. For CO_2_/HCO_3_^−^-buffered saline, NaCl was replaced by NaHCO_3_ and the solution was constantly aerated with 5% CO_2_ / 95% O_2_. The rate of change in intracellular H^+^ concentration was analyzed by determining the slope of a linear regression fit using OriginPro 8.6 (OriginLab Corporation). Recording and data analysis has been described in detail previously [84].

### Lactate imaging in cancer cells

Relative changes in intracellular lactate concentration in individual MCF-7 and MDA-MB-231 cells were measured using the lactate-sensitive FRET nanosensor *Laconic*. The specifications of the sensor have been described previously [86]. Transduction of MCF-7 and MDA-MB-231 cells with *Laconic* has been described in detail previously [29, 30]. In short, MCF-7 and MDA-MB-231 cells were plated on glass coverslips in serum-free Gibco OptiMEM medium (Life Technologies) and transduced with 4.8×10^10^ PFU of Ad5-Laconic (Vector Biolabs). After 4h RPMI medium was added and cells were incubated overnight under normoxic conditions. After 24h transduction was stopped by medium exchange and cells were incubated for at least three days under hypoxia (1% O_2_). Imaging experiments were performed in HEPES-buffered saline (143 mM NaCl, 5 mM KCl, 1 mM MgSO_4_, 1 mM Na_2_HPO_4_, 10 mM HEPES, 2 mM CaCl_2_, pH 7.2) in the nominal absence of CO_2_/HCO_3_^−^ at 21°C. Perfusion rate was set to 2 ml/min. Image acquisition was carried out with a Leica SP2 inverse confocal laser scanning microscope (Leica Microsystems IR GmbH), equipped with a 20x dry objective (HC PL APO 20x/0.70 CS). *Laconic* was excited at 458 nM and the emission light split into a 460-500 nm (mTFP) and a 520-550 nm (Venus) fraction. Pictures were taken at a sampling rate of 0.2 Hz. For ratiometric imaging of lactate concentration, the 460-500 nm fraction was divided by the 520-550 nm fraction. Data analysis was carried out with ImageJ and OriginPro 8.6.

### Determination of proton and lactate production in cancer cells

Extracellular pH values of cultured cells were monitored using a SDR SensorDish^®^ Reader (PreSens Precision Sensing GmbH). MDA-MB-231 and MCF-7 cells, respectively were plated in a 24-well HydroDish^®^ HD24 cell culture dish at a density of 10^5^ cells/well. Every well of the HydroDish^®^ is equipped with a pH sensor to allow online monitoring of extracellular pH during incubation. Cells were grown for three days under hypoxic conditions. Afterwards culture medium was exchanged by 600 μl of new medium, containing interfering antibody or inhibitors. Cells were incubated for 24h and pH values were recorded every 5 minutes. After the 24h incubation 500 ml of the medium were removed from the culture plate and deproteinized using perchloric acid and KOH precipitation. Lactate concentrations were then determined using an enzymatic assay (L-Lactic Acid (L-Lactate) Assay Kit, Megazyme u.c.). Lactate concentrations were normalized to cell number in each individual well. Therefore the cells were fixed with 4% PFA in the HydroDish^®^ and nuclei stained with 10 μM Hoechst 33342 (H1399; Thermo Fisher) in PBS. For each well, five pictures were taken at random locations and the number of cells was automatically counted using the software ImageJ as described previously [66].

### Determination of cell proliferation

Proliferation of MCF-7 and MDA-MB-231 cells was determined by counting of stained nuclei as previously described (Noor et al., 2018). In brief, cells were plated in 24-well plates at a concentration of 10^4^ cells/ml in RPMI medium, supplemented with antibodies or inhibitors. After 0, 1, 2, and 3 days of incubation cells were fixed with 4% PFA in PBS and nuclei stained with 10 μM Hoechst 33342 (H1399; Thermo Fisher). For each well, five pictures were taken at random locations and the number of cells was automatically counted using the software ImageJ as described previously [66].

### Molecular docking of CD147 and CAIX

CD147 and CAIX were docked and modeled as described previously [49, 72]. Docking of the two proteins was performed manually using the interactive graphical program COOT to maximize interface interactions followed by energy minimization at the complex interface using Crystallography and NMR System program (CNS) [87, 88]. The PDBs used to generate the model were 3B5H for hCD147 [89] and 4ZAO for CAIX [90]. Complexes were further analyzed and figures were generated using the graphics software Pymol (Schrödinger LLC).

### Calculation and statistics

Statistical values are presented as means ± standard error of the mean. For calculation of significance in difference one way analysis of variance (ANOVA) was carried out, followed by means comparison using either Scheffé or Bonferroni test, depending on whether datasets show homogeneity of variance or not. Homogeneity of variance was assessed using Levene’s test. All tests were carried out using OriginPro 8.6. In the figures shown, a significance level of p≤0.05 is marked with *, p≤0.01 with ** and p≤0.001 with ***.

## Acknowledgements

We thank Heike Kanapin, Sandra Pfeiffer, Hans-Peter Schneider and Patrick Pattar for excellent technical assistance. The work was funded by the Deutsche Forschungsgemeinschaft (to H.M.B.; BE 4310/6-1), the Research Initiative BioComp (to H.M.B), and by stipends from the Lotto-Stiftung Rheinland-Pfalz and the FAZIT Stiftung (to S.A.).

